# Molecular organization and mechanics of single vimentin filaments revealed by super-resolution imaging

**DOI:** 10.1101/2021.09.07.459174

**Authors:** Filipe Nunes Vicente, Mickael Lelek, Jean-Yves Tinevez, Quang D. Tran, Gerard Pehau-Arnaudet, Christophe Zimmer, Sandrine Etienne-Manneville, Gregory Giannone, Cécile Leduc

## Abstract

Intermediate filaments (IF) are involved in key cellular functions including polarization, migration, and protection against large deformations. These functions are related to their remarkable ability to extend without breaking, a capacity that should be determined by the molecular organization of subunits within filaments. However, this structure-mechanics relationship remains poorly understood at the molecular level. Here, using super-resolution microscopy (SRM), we show that vimentin filaments exhibit a ~49 nm axial repeat both in cells and *in vitro*. As unit-length-filaments (ULFs) were measured at ~59 nm, this demonstrates a partial overlap of ULFs during filament assembly. Using an SRM-compatible stretching device, we also provide evidence that the extensibility of vimentin is due to the unfolding of its subunits and not to their sliding, thus establishing a direct link between the structural organization and its mechanical properties. Overall, our results pave the way for future studies of IF assembly, mechanical and structural properties in cells.

## Introduction

The cytoskeleton of all metazoan cells is mainly composed of three types of filaments: microtubules, actin filaments (F-actin), and IFs. Cytoplasmic IFs, which exclude lamins localized inside the nucleus, form a network that extends from the nucleus to the plasma membrane and work in coordination with the other cytoskeletal filaments to control key cellular functions such as polarization, migration, division and signaling (*1–3*). IF networks also stabilize cellular architecture and protect cells from severe stretching (*4, 5*), confirming their mechanical role as cellular integrators described in the early literature (*6*). In particular, vimentin IF, a key marker of migrating mesenchymal cells, is essential for maintaining cell integrity and viability under conditions involving large deformations (*5*). This mechanical role was directly related to IFs interconnection with other cytoskeletal filaments and their synergetic ability to effectively disperse local deformations, allowing energy dissipation throughout the cell (*5*). The crucial role of IF is also highlighted by the fact that more than 90 human diseases are directly caused by mutations in IF proteins and that changes in IF composition also promote tumor growth and spreading (*7*).

Vimentin filaments exhibit assembly, structural and mechanical properties that strongly differ from F-actin and microtubules and that are also less understood at the molecular level. In terms of assembly, vimentin filaments result from the passive self-assembly of non-polar tetramers that associate laterally to form short filament precursors called unit-length-filaments (ULFs), which then fuse longitudinally during filament elongation (*7*). Simultaneously, vimentin also intercalate subunits along their length (*8–10*). Crosslinking experiments have shown that complex rearrangement of tetramers occurs during filament maturation (*11*), but structural information on vimentin at the level of filament is missing (*12*). Cryo-electron microscopy (cryo-EM) images showed that vimentin filaments consist of four protofibrils with a right handed supertwist and a helical pitch of ~96 nm, but the longitudinal positions of the tetramers within the protofibrils could not be distinguished (*13*). In terms of mechanical properties, vimentin filaments can be elongated up to 350% along their length without breaking in contrast with F-actin and microtubules that tend to break or disassemble at moderate strain (*14–17*). This high extensibility was attributed to the reversible and cooperative unfolding of alpha helices within vimentin subunits thanks to stochastic and numerical modeling (*14, 16*). Internal sliding of the subunits would also be consistent with these experimental observations, but the spatial resolution of the experiments was not sufficient to observe the exact molecular mechanism (*16*). Furthermore, most experiments designed to characterize the properties of vimentin were performed on purified proteins reconstituted *in vitro* and thus overlooked the potential role of cellular processes. For example, post-translational modifications occurring in the cytoplasm might modify the structural organization of IFs, and, as a result, impact their assembly and mechanical properties (*18*). Overall, the structural organization of vimentin filaments and their structure-mechanics relationship remain unclear both *in vitro* and within cells, primarily due to the lack of experimental techniques to resolve the molecular organization of proteins and to investigate their response to forces.

By delivering images with spatial resolutions below the diffraction limit of light, super-resolution microscopy (SRM) techniques have created new opportunities to study the nanoscale architecture of cellular structures directly in cells (*19, 20*). We used two major classes of SRM: (i) stochastic approaches based on Single Molecule Localization Microscopy (SMLM) such as PALM (*21*), STORM (*22, 23*), or DNA-PAINT(*24*) which typically achieve the best resolution; and (ii) selective deactivation of fluorophores by stimulated emission, minimizing the cross-sectional area of the focal spot below the diffraction limit such as STED (*25*). SMLM-based techniques allow to decipher for example the structural organization of integrin-based adhesions (*26, 27*) or nuclear pore complexes (*28*) at the nanoscale. The recent development of a cell-stretching device compatible with SRM methods, including SMLM, enables the exploration of protein deformation or reorganization upon mechanical strain (*29*). This technique also enables the study of the molecular mechanisms involved in mechano-sensitivity directly inside cellular structures (*29*).

In this manuscript, we used SRM techniques to reveal the organization of vimentin filaments at the molecular scale directly inside cells and *in vitro*, and bring new important insights into the IF assembly mechanism. Furthermore, by combining IF stretching with SRM, we showed the impact on external forces on the molecular organization of the filaments and unveiled their structure-mechanics relationship. This established a novel strategy to assess the tension applied on vimentin filaments in cells by quantifying their structural organization. Altogether, our results open the way to a better understanding of the structural, assembly and mechanical properties of IF in cells

## Results

### Vimentin ULFs are made of parallel tetramers inside cells

To probe the molecular organization of vimentin filaments directly inside cells, we first studied the early stage of filament assembly and focused on vimentin ULFs, using direct stochastic optical reconstruction microscopy (dSTORM)(*23*). ULFs are known to result from the lateral association of 8 to 10 tetramers, which are themselves made of the antiparallel association of parallel dimers (Fig. 1A) (*7*). Crosslinking experiments have shown that ULFs reconstituted *in vitro* are predominantly made of A11 tetramers in which there is an N-terminal overlap of the two antiparallel dimers (Fig. 1A) (*11*). The alternative types of tetramers - A22 in which there is an C-terminal overlap of the two antiparallel dimers, and A12 made of in-register antiparallel dimers (Fig. S1A) - were only found in mature filaments (*11*). However, whether the ULF organization is similar in cells or whether the cellular environment alters IF assembly has not been reported. To study ULFs in cells, we took advantage of the vimentin mutant vimentin-Y117L, which impairs longitudinal end-to-end annealing of filaments and thus stops filament assembly at the ULF stage (*30, 31*). The ULFs have previously been shown to form particles in cells and exchange subunits with the soluble tetrameric pool in an ATP-dependent manner (*32*). Vimentin-Y117L-GFP was expressed in mouse embryonic fibroblasts (MEF) vimentin-KO completely depleted of cytoplasmic intermediate filaments (*30, 31*). The GFP-terminal ends were labeled with GFP-nanobodies-AF647, which are fluorescently-labeled single-chain antibody domains with high affinity for GFP that are widely used for SRM (*33*). With the absence of unlabeled endogenous vimentin, the labeling of vimentin was maximized allowing to perform quantitative dSTORM imaging in optimal conditions (*23, 34*). We performed 2D-dSTORM imaging of fixed cells (Fig. 1B and Fig. S2A). While only large dots were observed in the GFP channel with epi-fluorescence, 2D-dSTORM images showed that the vimentin C-terminal ends formed doublets within the ULFs, consistent with ULFs being made of tetramers organized in a parallel fashion in cells. We estimated the spatial resolution of the images by measuring the full width at half maximum of all the doublet dots and found 27 nm on average (s.d. 10 nm). The distance between the C-terminal ends, hereafter called pairwise distance, was quantified using a double Gaussian fit (Fig. 1C) and found to be ~50 nm on average (s.d. 14 nm)(Fig. 1D). This distance is significantly smaller than the ULF length of ~60 nm measured *in vitro* using electron microscopy (*35, 36*).

**Fig. 1:**
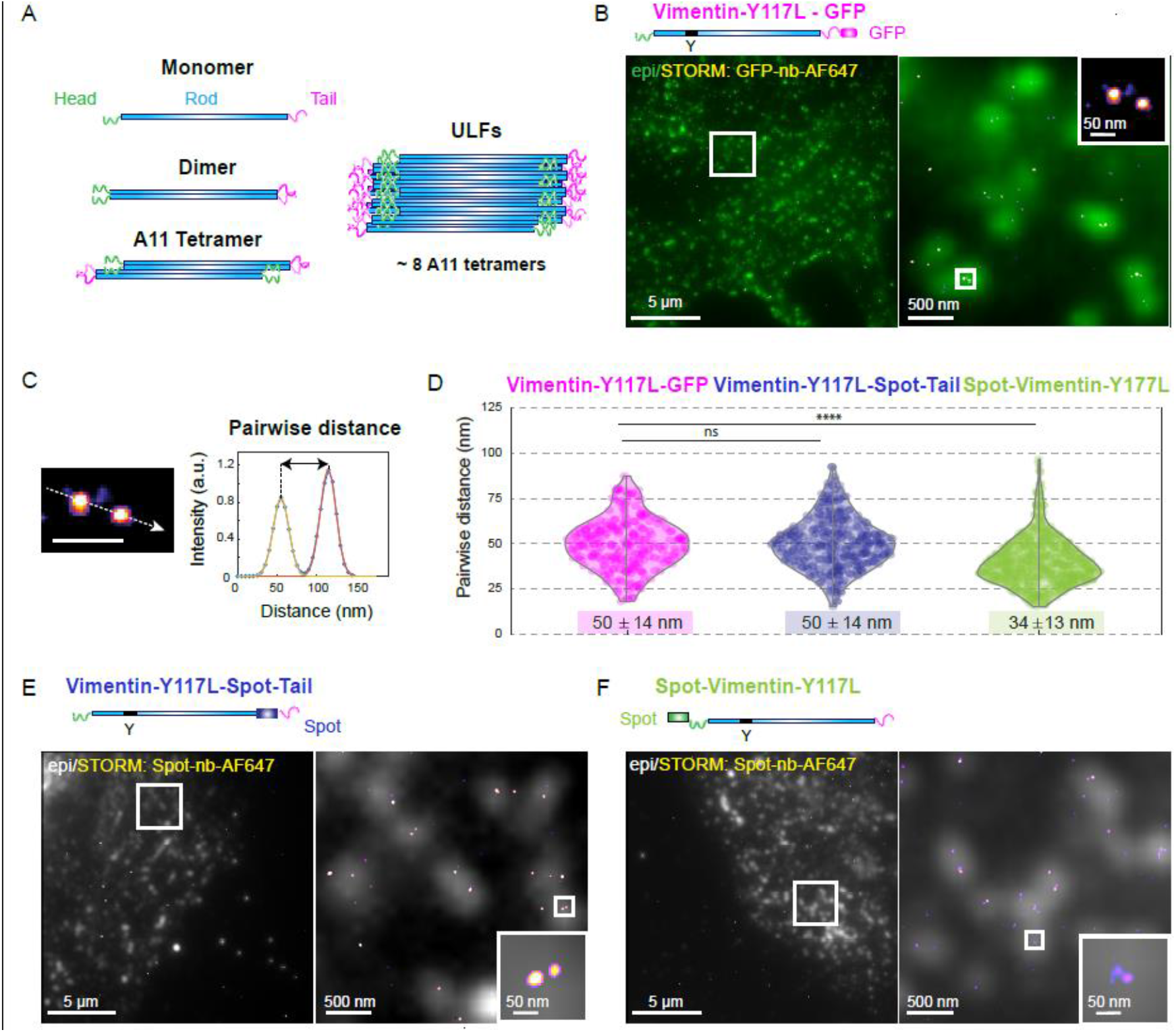
Structural organization of vimentin ULFs in cells. **(A)** Schematic of vimentin assembly. Vimentin monomers are formed by a head (green), a rod (blue) and a tail (purple). Two parallel dimers associate to form anti-parallel, half-staggered tetramers. About 8 tetramers associate laterally to form vimentin precursors called ULFs. **(B)** Top: Schematic of the Vimentin-Y117L-GFP protein. Bottom: epi-fluorescence image and a higher magnification the boxed region of a MEF vimentin-KO cell expressing the vimentin-Y117L-GFP construct (green) superimposed on the corresponding 2D-dSTORM image (fire) acquired with GFP-nanobodies-AF647. Inset: magnifications of the boxed region. **(C)** Magnification of a ULF from the inset in (B). The fluorescence intensity profile along the ULF is fit by two Gaussian and pairwise distance is calculated from the distance between the two peaks. **(D)** Distributions of pairwise distances measured with the vimentin-Y117L-GFP (magenta), vimentin-Y117L-Spot-Tail (blue) and Spot-vimentin-Y117L (green) constructs expressed in MEF vimentin-KO cells. Mean values ± standard deviations of the total distributions are written below the violin plots. Data represent 253, 317 and 596 ULFs respectively, collected from 9, 10 and 16 cells from 3 to 4 independent experiments. P value was calculated with a nested *t*-test of the 3 repeats, ****: *P* < 0.0001, n.s.: non-significant. **(E-F)** Top: Schematic of the Vimentin-Y117L-Spot-Tail protein (E) and Spot-Vimentin-Y117L protein (F). Bottom: epi-fluorescence image (grey) and a higher magnification the boxed region superimposed on the corresponding 2D-dSTORM image (fire) of Spot-nanobodies-AF647 (fire). Inset: magnifications of the boxed regions.

To understand the discrepancy between the ULF length measured *in vitro* with EM and in cells, we first investigated whether this difference was due to the GFP location or size. Vimentin proteins are divided in three parts: a central alpha-helical rod domain flanked by two non-alpha-helical end domains, a head in N-terminal and a tail in C-terminal (Fig.1A). Because the GFP tag is localized at the end of the disordered tail domain, its relative position to the end of the rod domain is not clearly defined. Vimentin tails may swivel around the end of the rod domain and thereby impact the measurements of the distance between the C-terminal ends. To test this hypothesis, we reduced the size of the tag by replacing the GFP by a 12 amino acid tag, called “Spot” (*37*), inserted between the rod and the tail (Fig. 1E). We expressed the vimentin-Y117L-Spot-Tail construct in vimentin KO MEF cells, fixed them and performed immuno-labeling with AF647-labeled Spot-nanobodies to perform 2D-dSTORM images (Fig. 1E). As found with GFP, ULFs also appeared as dots in epi-fluorescence images and doublets in 2D-dSTORM images, with an average length of ~50 nm (Fig. 1D). The absence of a significant difference for doublet size measured with C-terminal GFP-tagged vimentin and vimentin with a Spot-tag between the rod and tail confirms that probe size and vimentin tail have no impact on ULF length measurement.

To further investigate the length discrepancy, we tested whether other types of tetramers than A11, such as A22 and A12 (Fig. S1A) could form in cells. To do so, we quantified the localization of vimentin N-terminal heads within ULFs (Fig. 1A). We designed a Spot-vimentin-Y117L construct that was expressed in MEF vimentin KO cells and performed 2D-dSTORM imaging (Fig. 1F). We observed that vimentin heads were also organized in doublets within ULFs, but we measured a significantly smaller doublet length of ~34 (s.d. 13 nm) compared to C-terminal ends. These results are consistent with an organization of vimentin tetramers with C-terminal ends on the outside and N-terminal ends on the inside of ULFs, and thus indicate that ULFs are predominantly composed of A11 tetramers in cells. Importantly, we confirmed the predominance of A11-type over A22 and A12 structural organization by simultaneously observing vimentin heads (labeled by a Spot-tag) and tails (labeled by a GFP) using dual color STED imaging of Spot-nanobodies-AF594 and GFP-nanobodies-Star635 (Fig. S1B).

### Vimentin ULFs are ~59 nm long and are tilted in cells

We then investigated whether the discrepancy between ULF length measured *in vitro* with EM and in cells with 2D-dSTORM could result from tilted ULFs projected in 2D. We used a 3D-dSTORM microscope equipped with two objectives and two cylindrical lenses which allows for improved spatial resolution in all three dimensions (< 25 nm in each direction), compared to a single cylindrical lens combined with one objective (*38*). These experiments showed that ULFs were tilted in cells. The vimentin tails formed doublets, whose orientation could be quantified with an average angle of ~22° (s.d. 17°) relative to the horizontal, a value which is underestimated since the population of vertical ULFs could not detected with our analysis method (Fig. 2 A-C). Two populations of orientation were observed: ~46% of ULFs had an angle of less than 15° (Fig. 2C), which could correspond to the population of ULFs aligned with the microtubule network almost parallel to the plasma membrane (*39*) (Fig. S3) and the other population, with a higher tilt value, which could then correspond to “isolated” ULFs with a random orientation. To test this hypothesis, we quantified the ULF length distribution and orientation after microtubule disassembly induced by nocodazole treatment. While the ULF orientation became more uniformly distributed compared to untreated control cells (Fig. 2D), neither the mean ULF 3D length nor the 3D width of the distribution were significantly altered by microtubule depolymerization (Fig. 2D). The 2D projected doublet size was significantly shorter than in control (Fig. 2D), as observed in 2D-dSTORM experiments (Fig. S4), due to the loss of ULF orientation after microtubule depolymerization. These results also indicate that microtubule-associated motors do not alter the length of ULFs in cells. Inhibition of myosin II activity by Blebbistatin did not impact ULF length distribution in 2D (Fig. S4), suggesting that actin-associated motors do not alter ULF orientation and length. Taking into account the 3D coordinates, the doublet size was measured to be on average 59 nm (s.d. 15 nm) (Fig. 2D), in agreement with ULF length measured *in vitro* (*35, 36*). We then corrected the distance between N-terminal ends for projection error and found a distance of ~39 nm. Our results unambiguously demonstrate that the length and structural organization of cellular ULFs, made up mostly of A11 tetramers, are similar to the ones reported in reconstituted systems (*11, 35, 36*) and that the orientation of ULFs in cells is determined by their interaction with the microtubule network, but not their length.

**Fig. 2:**
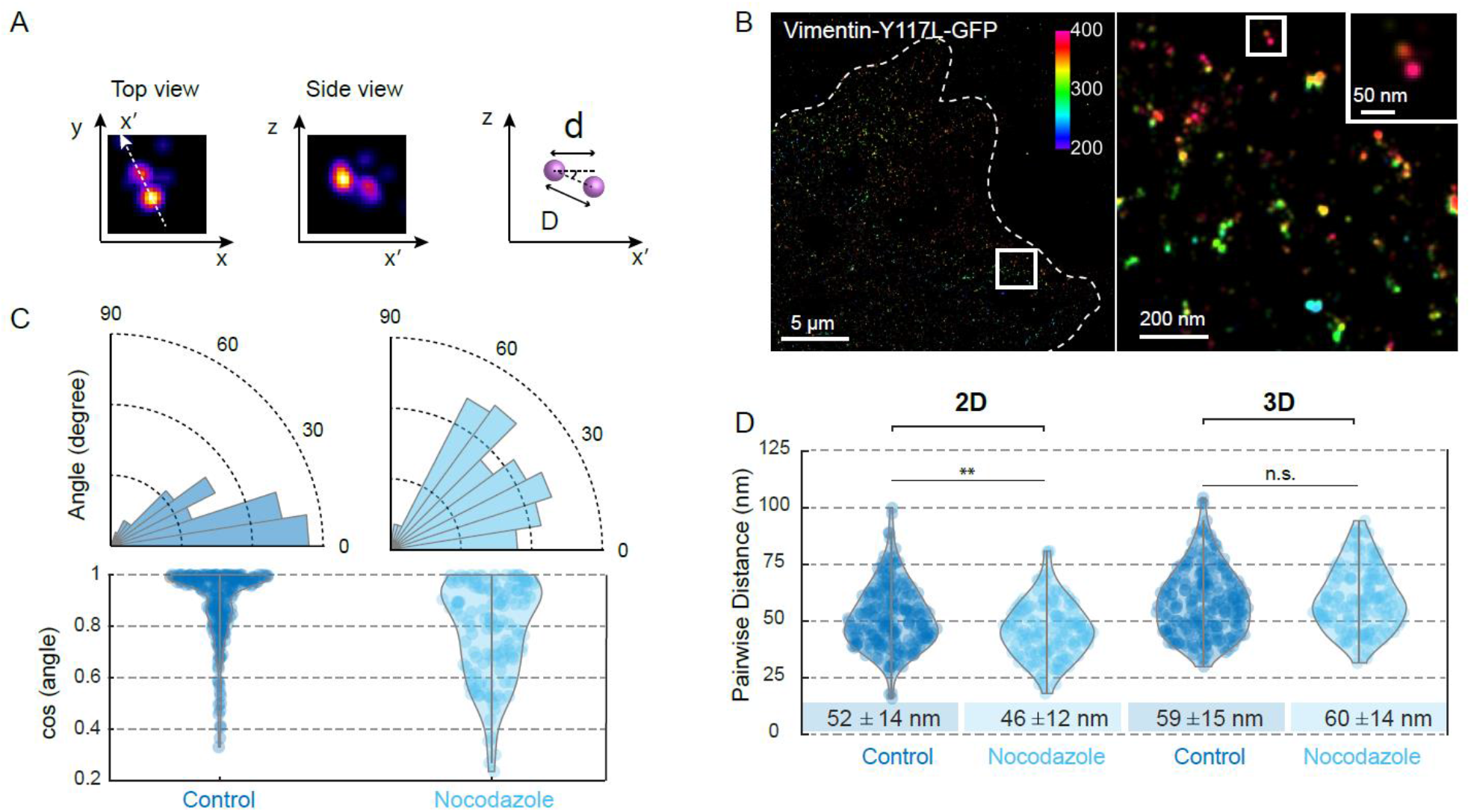
Vimentin ULFs are ~59 nm long and are tilted in cells. **(A)** Top and side views of a ULF imaged by 3D-dSTORM, and a cartoon showing the projected distance “d” measured in 2D-dSTORM, the real distance “D” between vimentin-Y11L-GFP C-terminal ends in ULFs measured in 3D, and the tilt angle. **(B)** 3D-dSTORM image and a higher magnification the boxed region of a MEF vimentin KO cell expressing vimentin-Y117L-GFP and labeled with GFP-nanobodies-AF647. The color indicates the depth *z* in nm. Inset: magnification of the boxed region showing a ULF. Images were acquired using a dual-objective microscopy setup with cylindrical lenses (see methods). **(C)** Distributions of ULFs angles relative to the substrate and cosinus of ULF angles in control cells and in cells treated with nocodazole. **(D)** Distributions of pairwise distances between vimentin-Y11L-GFP C-terminal ends in ULFs projected in 2D (distance “d”) and measured in 3D (distance “D”) in control and nocodazole treated MEF vimentin-KO cells. Mean values ± standard deviations of the total distributions are written below the violin plots. 204 and 133 ULFs were analyzed from 4 control cells and 5 cells treated with nocodazole respectively, acquired during 2 independent experiments. *t*-test, **: *P* < 0.01, n.s.: non-significant.

### Vimentin ULF length is heterogeneous in cells

The distribution of pairwise distances between C-terminal tails, which we use as an estimate of ULF length, is broad around the mean value of 59 nm with a standard deviation of 15 nm, i.e. ~25% of the mean length. This broad distribution could either result from uncertainties in the SRM measurements or could be intrinsically associated with the molecular organization of the ULFs, since we ruled out the impact of the tail, the probe size and position as well as of any previous stretching by molecular motors (Fig. 1D-E and 2D). To quantify the possible errors coming from SRM measurements, we measured the nuclear pores radius, which has become a standard for quantitative SRM (*34*), using a similar labeling technique (imaging of Nup96-SNAP labeled with SNAP-AF647), the same 3D-dSTORM microscopy setup and quantification method (Fig. S5). We measured a nuclear pore radius of 58 nm (s.d. 6 nm) with a narrower distribution, indicating that the width of the distribution for vimentin ULF lengths is not inherent to the imaging technique. Moreover, we did not observe any correlation between ULF length and the mean peak intensities of the doublet (Pearson correlation coefficient of 0.1, P value for the non-zero correlation hypothesis of 0.3), suggesting that the length heterogeneity is not related to fluctuations in the number of tetramers forming the ULF (Fig. S6). Altogether, these results show that the length of vimentin ULFs is intrinsically heterogeneous in cells, which may impact the regularity of filament organization after assembly.

### The organization of vimentin tails exhibits an axial repeat of 49 nm within filaments in cells

The internal organization of mature vimentin filaments is still poorly understood, since the exact organization of tetramers within the filaments is unclear (*12*). Indeed, it was previously shown in biochemical studies that while ULFs consist mainly of A11-type tetramers, mature vimentin filaments also contain A22-type and A12-type tetramers in addition to the A11 (Fig. S1A), suggesting that major rearrangements of tetramers occur during longitudinal annealing of filaments (*11*). Since the commonly accepted mechanism for filament assembly is the fusion of vimentin ULFs, we investigated whether we could observe a signature of such organization by quantifying the position of the head and tail domains in mature filaments using super-resolution microscopy. We first used a vimentin construct with a C-terminal GFP expressed in MEF vimentin KO cells and labeled with GFP-nanobodies-AF647, and performed 2D-dSTORM images of vimentin filaments in the TIRF mode (Fig. 3A). We clearly observed an axial repeat along the filaments, reflecting the organization of the vimentin tails. We developed an image analysis software (see methods), that allows automatic detection of intensity peaks and measurement of the pairwise distance between peaks along the filaments (Fig. 3B-C). The distribution of pairwise distances between the vimentin tails shows a maximum at ~ 49 nm, which we take as an estimate of the axial repeat (Fig. 3D). Because the observed filaments were a few microns long and localized in a layer of less than 100 nm above the surface in the TIRF mode, their tilt angle was less than 5° and the error on the axial repeat estimation was less than 0.4%. The 3D-dSTORM images also confirmed that most filaments were parallel to the glass surface (Fig. S7). We also verified that this organization was not cell specific and that cell fixation had no impact on the axial repeat measurements and nor did microtubule depolymerization (Fig. S8–9). SRM reveals the structural organization of mature vimentin filaments which display a ~49 nm axial repeat of the tails, demonstrating that the filaments are made up of a succession of ULFs (Fig. 3E).

**Fig. 3:**
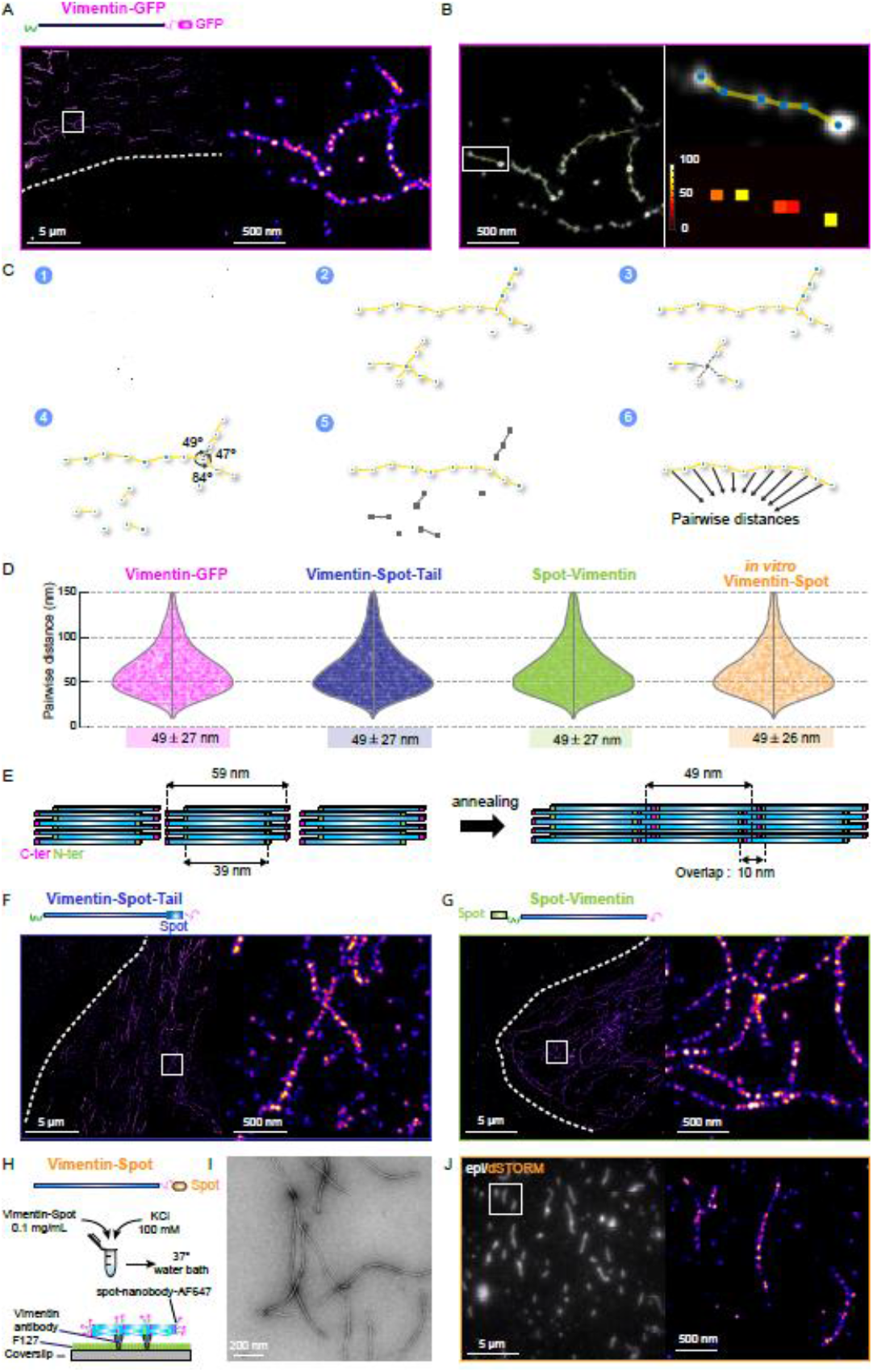
Structural organization of vimentin filaments in cells and filaments reconstituted *in vitro*. **(A)** 2D-dSTORM image and a higher magnification of the boxed region of a MEF vimentin-KO cell expressing vimentin-GFP and labeled with GFP-nanobodies-AF647. Dashed line indicates the cell edge. **(B)** Schematics of a filament and how pairwise distances are measured. Left: 2D-dSTORM image from (A) superimposed with intensity peaks which are automatically detected (labeled in blue) and with the distances to the closest neighbors depicted in yellow (see methods). Right: Zoom of the boxed area and color map of the pairwise distances. **(C)** Schematics of the analysis method for the pairwise distance measurements (see methods). **(D)** Distributions of pairwise distances between vimentin-GFP tails (magenta), vimentin-Spot-Tails (blue), Spot-vimentin heads (green) measured in cells and vimentin-Spot tails (yellow) measured in *in vitro* reconstituted filaments. 4664, 13936 and 15903 pairwise distances were measured respectively from 10 cells per condition acquired in each case in 3 independent experiments, and 4167 pairwise distances were quantified from 3 independent experiments for reconstituted filaments. The distribution modes and standard deviations are written below the violin plots. **(E)** Schematic of the intertwining of ULFs after end-to-end annealing. **(F-G)** 2D-dSTORM image and a higher magnification of the boxed region of MEF vimentin-KO cells expressing vimentin-Spot-Tail (F) and Spot-vimentin (G) and labeled with Spot-nanobodies-AF647. Dashed line indicates cell edges. **(H)** Protocol of vimentin-Spot filament assembly. After dialysis in a 2.5 mM phosphate buffer, vimentin assembly is started by addition of KCl (final concentration of 100 mM) to vimentin tetramers (0.1 mg/mL) and incubation for 3h at 37°C. **(I)** Representative TEM image of vimentin-Spot filaments assembled *in vitro*. **(J)** Epi-fluorescence images (grey) superimposed on the corresponding 2D-dSTORM images (fire) of vimentin-Spot filaments. Right: 2D-dSTORM image of the boxed region.

Next, we tested whether the size and localization of the GFP tag could impact the estimation of the axial repeat along the filaments by 2D-dSTORM of a vimentin-Spot-Tail construct (Fig. 3F). The distribution of pairwise distances showed no significant difference compared with that obtained with vimentin-GFP, showing that neither the tag size nor tail swiveling had any impact on the axial repeat (Fig. 3D). Because the vimentin tail has also been implicated in vimentin organization and remodeling in many cell types (*2*), we tested whether tail-less vimentin displayed similar organization using a construct vimentin-ΔTail-Spot, but we also did not find any significant difference in the axial repeat with vimentin-GFP (Fig. S9 A-B). Finally, we also showed that a similar axial repeat of vimentin tails was present in endogenous IF network in astrocytoma cells after 2D-dSTORM imaging of vimentin tails, using site specific labeling via a monoclonal antibody targeting an epitope on the tails (clone V9) (Fig. S9 C-D). This indicates that the characterized structural organization of vimentin is not an artefact due to overexpression of vimentin proteins. These results show that the organization of vimentin tails display an axial repeat of ~49 nm along the filaments in cells, which is smaller than the 59 nm measured in ULFs (Fig. 2D). This is quantitatively consistent with a scenario where ULFs partially overlap by ~10 nm during the end-to-end annealing process of filament assembly and without major structural reorganization of the ULFs (Fig. 3E). If the spatial resolution of the system was less than 10 nm, we would expect an alternation of 39 and 10 nm of the tails due to this partial overlap of ULFs (Fig. 3E). With a resolution of ~27 nm, the tails of successive overlapping ULFs ends (10 nm apart) appear as single spots, leading to an axial repeat of 49 nm.

### Vimentin filaments intertwine during the fusion process leading to filament assembly

To further validate this scenario of ULF intertwining, we investigated the organization of vimentin heads within filaments in cells using the N-terminal Spot-tag and 2D-dSTORM imaging of Spot-nanobodies-AF647 (Fig. 3G). We observed a distribution of pairwise distances similar to that of vimentin tails with a main peak also at ~49 nm (Fig. 3D). Although longer than the distance between vimentin heads in ULFs that we estimated to be ~39 nm (Fig.2D and 3E), this axial repeat is still compatible with the picture that ULFs partially overlap during the annealing process. Indeed, in principle, we would expect an alternation of 39 and 10 nm for the N-terminal heads (Fig. 3E), but the SRM setup does not have the spatial resolution to resolve this pattern. As in the case of vimentin tails, the heads of consecutive overlapping ULF ends appear as single spots leading to a global axial repeat of 49 nm. Overall, our results allow us to provide strong evidence of ULFs intertwining during filament assembly and to estimate of the overlap distance of ULFs directly inside cells.

### Vimentin structural organization is similar in *in vitro* reconstituted filaments and in cells

Since previous studies on vimentin have been performed with filaments reconstituted *in vitro*, we tested whether the filament organization we found in cells was similar *in vitro*. We designed a vimentin construct with a Spot-tag at the C-terminal end and purified it from bacteria following a previously described protocol (*40*). The reconstituted filaments were assembled by addition of salt (Fig. 3H). Using electron microscopy (EM), the Spot-tag was checked to have no impact on the formation of filaments (Fig. 3I). Then, we performed 2D-dSTORM imaging of non-fixed filaments attached on a glass surface via anti-vimentin antibodies and immuno-stained with Spot-nanobodies-AF647 (Fig. 3H-J). The distribution of pairwise distances between vimentin tails displayed an axial repeat of ~49 nm and did not exhibit significant differences with that observed inside the cell (Fig. 3D). We also verified that *in vitro* ULFs, obtained after 2 second of assembly, had a similar structural organization and length than in cells (Fig. S2B). Altogether these results show that the internal organization of filaments *in vitro* and inside the cell are similar, suggesting that interactions with cellular proteins have a limited impact on vimentin architecture. The 49 nm axial repeat of the tails can therefore be used as a “ruler” for vimentin filaments.

### Mechanical forces exerted on reconstituted filaments increase the axial repeat

Vimentin filaments have remarkable mechanical properties as they can be stretched *in vitro* (*14–17*). Two molecular mechanisms are in principle consistent with the extensibility of single filaments: either unfolding of the alpha-helices or the sliding of tetramers within the filaments (*14, 16*). We studied the impact of single filament stretching on the axial repeat to decipher which molecular mechanism is involved. We used a stretching device compatible with all the SRM techniques that was used to study mechano-sensing in cells (*29*). We directly attached the reconstituted filaments to the stretchable PDMS substrate using anti-vimentin antibodies, as on glass, and applied a uniaxial stretching step of 30% amplitude which increased the contour-length of the filaments by ~18% (averaged for all the directions) (Fig. 4A-B). We performed 2D-dSTORM images of vimentin tails labeled by Spot-nanobodies AF647 without or with stretching (Fig. 4B) and computed the pairwise distances parallel (0 ± 10°) and perpendicular (90 ± 10°) to the stretch axis. We observed a significant increase in pairwise distances parallel to the stretching axis compared to perpendicular distances, which was not present in the control experiment performed on the stretching device but without stretching (Fig. 4C-E). Although the frequency of pairwise distances around 64 nm, corresponding to a 30% increase compared to the case without stretching, was higher in the parallel case (Fig. 4D), the maximum of the distribution was shifted by only ~5 nm, suggesting that not all subunits are unfolded during stretching. Our results thus provide direct evidence that vimentin subunits change conformation during stretching, and also give fundamental insights into the link between molecular architecture and mechanical properties of single filaments. Additionally, these results also reveal that pulling on filaments induces an increased distance between vimentin tails, indicating that the axial repeat length may be taken as a readout of the tension applied on the filaments.

**Fig. 4:**
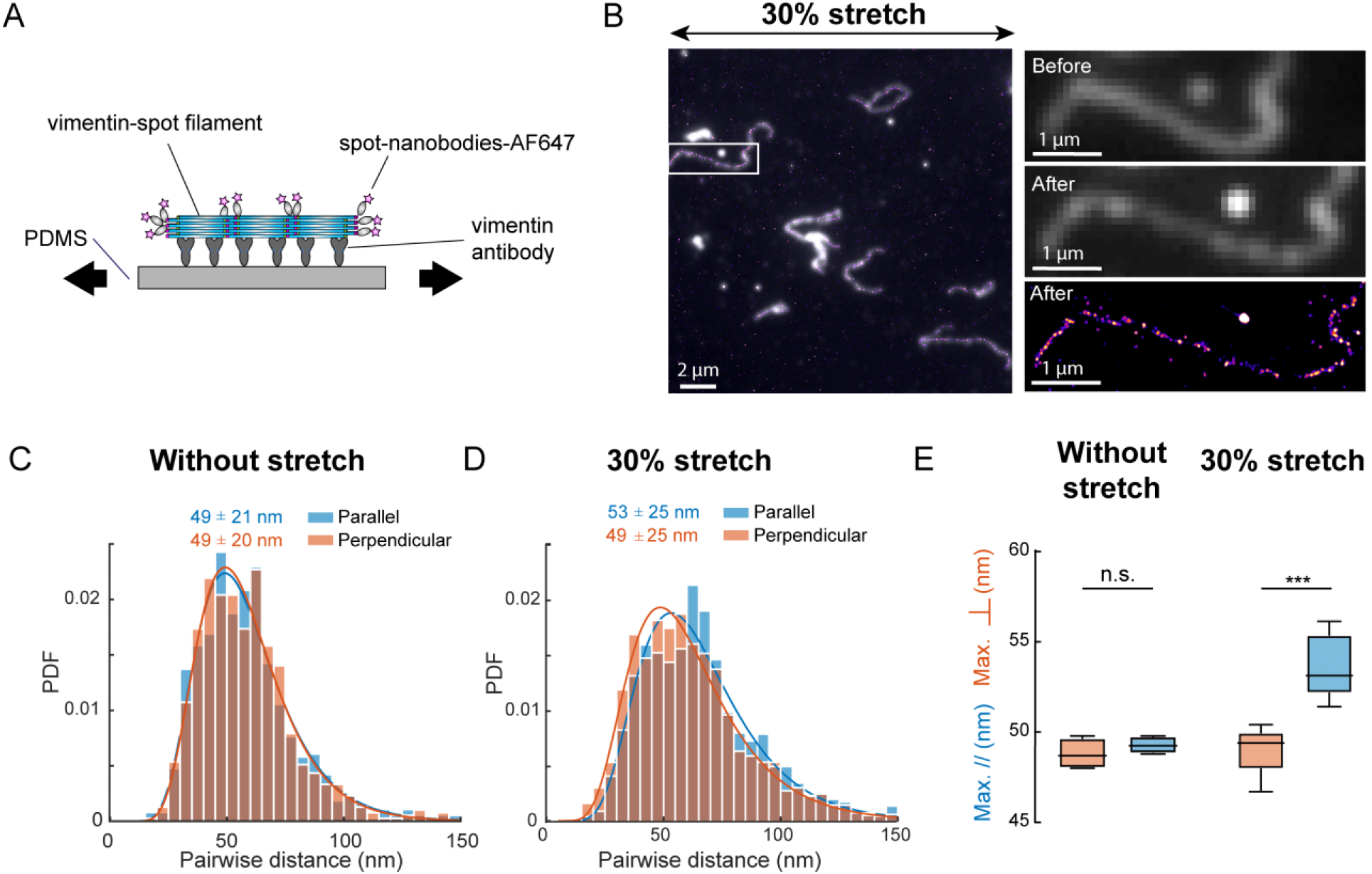
Structural organization of stretched vimentin filaments reconstituted *in vitro*. **(A)** Schematic of vimentin-Spot filament attachment to the PDMS subtract through vimentin antibodies and immune-labeled by Spot-nanobodies-AF647. The subtract is passivated by BSA. **(B)** Epi-fluorescence images (grey) superimposed on the corresponding 2D-dSTORM images (fire) of vimentin-Spot filaments stretched 30%. Right: Higher magnification of the boxed region before and after stretching (epi and dSTORM). **(C-D)** Histograms of pairwise distances aligned with the uniaxial stretch (parallel, blue) and perpendicular to the stretch direction (perpendicular, orange) before stretch (C) and for a stretch of 30% (D). PDF stands for probability density function. Blue and orange curves correspond to a lognormal fit. The distribution modes and standard deviations are written above the curves. 753 parallel and 779 perpendicular pairwise distances were measured before stretching and 1755 parallel and 1452 perpendicular pairwise distances were measured after 30% stretch, acquired from 3 independent experiments. No significant difference was observed between the parallel and the perpendicular distributions before stretching and a **** difference was observed for the 30% stretch condition (*P* < 0.0001). **(E)** Box and whiskers (min to max) of the modes of the lognormal fit of the pairwise distances parallel (blue) and perpendicular (orange) to the stretch direction. 4 dSTORM images were analyzed before stretch and 5 after 30% stretch, from 3 independent experiments. Paired *t*-test was used to compare the two conditions. ***: *P* < 0.001

### Vimentin filaments are not elongated after cell stretching up to 50%

Finally, we studied the impact of cell stretching on the axial repeat directly in cells. We applied a step of uniaxial stretch up to 50% amplitude to live cells using the stretching device compatible with SRM, followed by immediate fixation of the cell after having reached the stretching plateau (in less than 1 min). First, we used vimentin-KO MEF cells expressing vimentin-GFP and plated them on the fibronectin-coated stretchable PDMS substrate. Low resolution epi-fluorescence images of vimentin-GFP allowed us to observe the reorganization of the vimentin network axis before and after stretching (Fig. 5A). Comparison of the mean directionality of the filaments before and after having reached the stretching plateau showed that the filaments reoriented along the stretching axis, attesting that external forces were transmitted to the filaments (Fig. 5B). We also quantified the extension of single filaments before and after stretching and observed that the length increase was less than 6 % on average along the stretch axis and for a stretch amplitude of up to 50% (Fig. 5C). Next, we analyzed the structural organization of vimentin filaments using DNA-PAINT with a HALO-tag at the C-terminal end of vimentin, as previously described (*29*). The DNA-PAINT method uses the stochastic binding of a fluorescent ligand to localize the target protein. DNA-PAINT is based on hybridization of complementary DNA strands, one located on the target protein (docking strand) and the other one on the dye (imager strand) (*24*). We switched from dSTORM of vimentin-GFP to DNA-PAINT with vimentin-HALO, to further improve the quality of the images. We analyzed the pairwise distance between vimentin tails and quantified the axial repeat with or without stretching. Comparison between pairwise distances parallel (0±10°) and perpendicular (90±10°) to the stress axis did not show any significant difference for 35 % and 50% stretch, and no difference was observed with control cells that were not stretched (Fig. 5D-G, Fig. S10). The lack of significant filament extension or change in the internal organization of vimentin within the filaments indicates that cytoplasmic vimentin filaments are not under tension inside the cells, even after having reached a 50% stretch, or that they quickly release the applied tension.

**Fig. 5:**
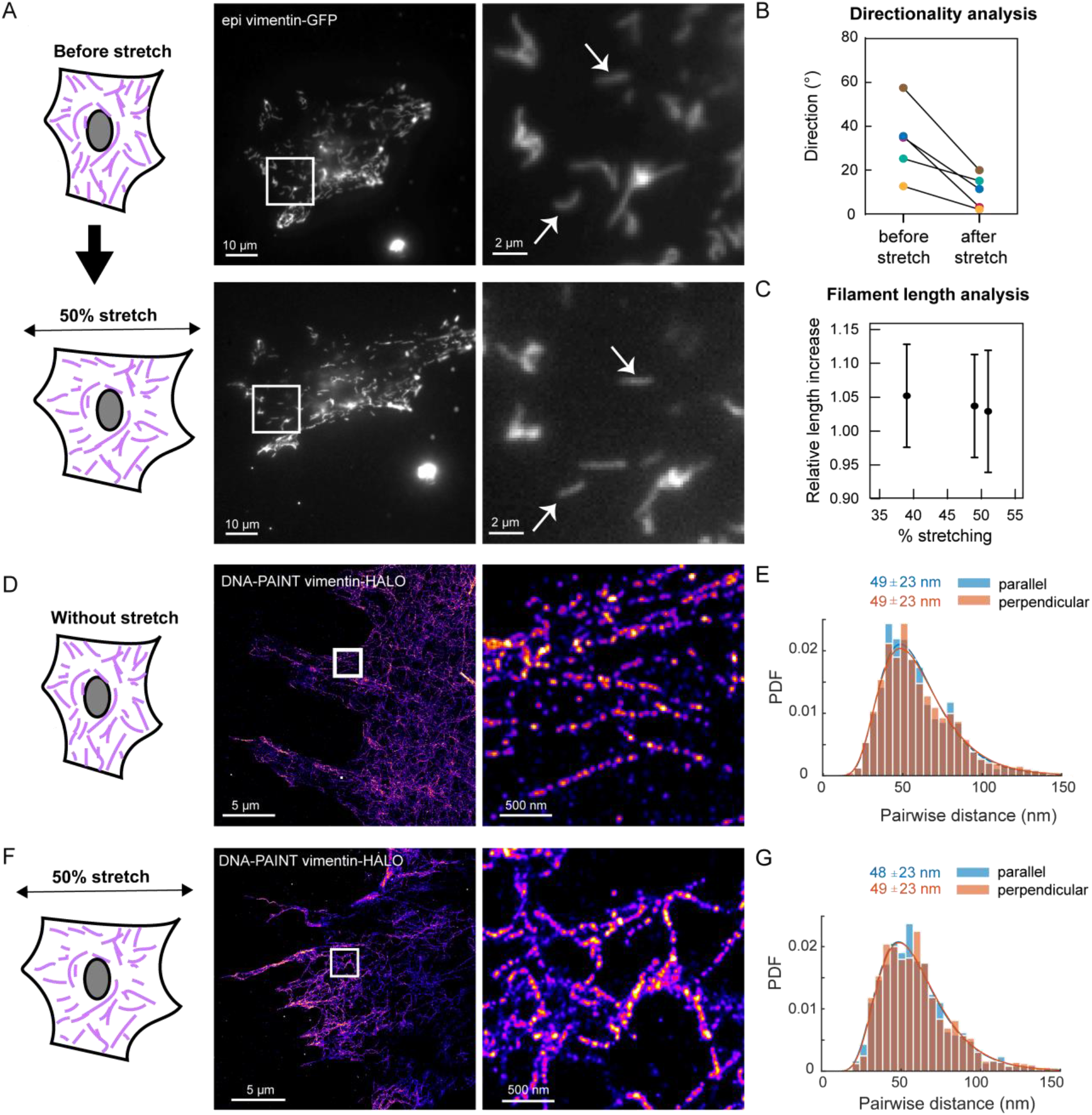
Structural organization of vimentin filaments within stretched cells. **(A)** Epi-fluorescence images of a MEF vimentin-KO cell expressing vimentin-GFP before and after 50% uniaxial stretch and higher magnification of the box regions. White arrows: filaments which are not stretched in the cell **(B)** Directionality analysis performed using the Fiji plugin was performed before and after stretching of 5 cells. 0° corresponds to the direction parallel to the stretch axis. **(C)** Relative length increase measured in 3 live cells after 39 to 51% stretch. **(D)** DNA-PAINT image of a MEF vimentin-KO expressing vimentin-HALO and stretched 50%. Middle: magnification of the boxed region. Right: corresponding map of pairwise distances. **(E)** Histogram of pairwise distances aligned with the uniaxial stretch (parallel i.e. with an angle of 0 ± 10°, blue) and perpendicular to the stretch direction (perpendicular i.e. with an angle of 90 ± 10°, orange), detected from the DNA-PAINT image in D (stretch 50%). Blue and orange curves correspond to a lognormal fit. The distribution modes and standard deviations are written above the curves. 3823 and 4157 pairwise distances were measured respectively from 2 independent experiments. No significant difference was observed between the parallel and perpendicular distributions. **(F)** Left: DNA-PAINT image of a MEF vimentin-KO expressing vimentin-HALO and not stretched. Middle: magnification of the boxed region. Right: corresponding map of pairwise distances. **(G)** Histogram of pairwise distances aligned with the uniaxial stretch (parallel, blue) and perpendicular to the stretch direction (perpendicular, orange) in a control cell without stretching (stretch 0%). Blue and orange curves correspond to a lognormal fit. The distribution modes and standard deviations are written above the curves. 1242 and 1016 pairwise distances were measured respectively, from 2 independent experiments. No significant difference was observed between the parallel and the perpendicular distributions. PDF stands for probability density function.

## Discussion

Although vimentin IFs play a crucial role in many key cellular functions such as mechanical integrity, polarization, invasion and mechano-sensing, our understanding of their structural organization is much more limited compared to F-actin or microtubules. Here, using site-specific labeling and SRM, we unveiled the architecture of vimentin filaments at the molecular level directly inside the cell and in reconstituted filaments *in vitro*. We showed that vimentin ULFs are mostly composed of A11 tetramers with a mean length of ~59 nm with a broadly distributed size in cells. We then analyzed the organization of vimentin proteins within mature filaments both in cells and in reconstituted filaments. In both cases, the filaments showed an axial repeat of ~49 nm in the organization of head and tail domains, consistent with a mechanism where the ends of vimentin ULFs intertwine during filament assembly. Finally, using a stretching device compatible with SRM, we probed the impact of external stretching on the structural organization of filaments both *in vitro* and within cells. In the case of purified filaments, the axial repeat was increased for stretching amplitudes up to 30%, only in filaments parallel to the stretch axis. This indicates that mechanical strain leads to the unfolding of alpha helices in vimentin subunits. Interestingly, the axial repeat of intracellular filaments and their contour length did not increase in cells stretched up to 50%. Instead, the filaments reoriented in the direction of the stretch, indicating that the filaments were not under tension or that they released it quickly.

Starting with vimentin ULFs, we observed using SRM that the distribution of distances between C-terminal ends was broad (s.d. 15 nm), reflecting the ULF polymorphism and size heterogeneity in cells. Because tetramers have been reported to be loosely packed within the ULFs (*41*), the size heterogeneity could also reflect a partial misalignment of the tetramers. It may also be related to the variety of the ULF types, which are mostly formed of A11 tetramers, but could also include A22 or A12 tetramers (Fig. S1B). In addition, using an improved 3D-dSTORM imaging with 25 nm resolution in all three dimensions, we showed that the 3D information should not be overlooked. Indeed, it provides crucial knowledge on the ULF orientation that may be important in the assembly process. One remaining question is whether the different types of ULFs could anneal during filament assembly and whether the type of ULF impacts the assembly rate. Polymorphism after IF assembly has previously been described in terms of tetramer types as well as the number of tetramers per unit length (*7*), but only for filaments reconstituted *in vitro*. Here, we go one step further by describing this polymorphism directly inside the cell. We observed that the axial repeat reflecting the succession of ULFs within filament is also heterogeneous, which suggests that ULFs of different sizes may anneal together. Improvement of live SRM techniques will be necessary to achieve a better spatial and temporal resolution in 3D, in order to study the dynamic assembly of filaments within cells and to understand the potential functions associated with this polymorphism. Moreover, resolving the sub-ULF organization within filaments will require the use of SRM technique having a resolution close to 1 nm such as MINFLUX (*42*).

Our results brought significant advances in the understanding of vimentin structural organization and assembly. A previous study based on theoretical modeling of crosslinking experiments estimated the size of the axial repeat to be 42.7 nm (*43*). However, there was no direct experimental measurement of this length. Using SRM, we showed that the size of the axial repeat is ~49 nm. As ULFs in cells measure 59 nm, this demonstrates a partial ULFs overlap (~10 nm) occurring during filament assembly (*40*). Thus, our results provide direct evidence of the ULF intertwining within filaments, validating previous theoretical predictions (*43*). Interestingly, the intertwining of ULFs does not hinder the addition and removal of subunits along the filaments as observed in cells and *in vitro* (*8–10*), suggesting that the bond between tetramers can be broken spontaneously. Of note, 49 nm is also half the helical pitch of vimentin filaments measured by cryo-EM (*13*), suggesting that the helical pitch is formed by two successive ULFs. However, we do not have the spatial resolution with SRM to resolve the protobribils within filaments, and measure the tilt orientation of these protofibrils with respect to the main axis of the filament. Cryo-EM and SRM therefore bring complementary descriptions of vimentin architecture. While cryo-EM resolve the protofibril organization and describes their helical path, SRM describes the size of the ULF after insertion within filaments regardless of the orientation of the protofibrils within the filament, and this is not accessible by cryo-EM since vimentin head and tails positions could not be distinguished with this technique. We also showed that vimentin filaments in cells and reconstituted *in vitro* displayed similar axial repeat, suggesting that the molecular organization of IFs is similar, and thus providing further validation of the physiological relevance of the bottom up approach (*12*). Furthermore, because IF assembly and disassembly are regulated by phosphorylation and other post-translational modifications (*18*), it will be interesting to investigate whether these modifications alter the axial repeat of IFs, reflecting changes in their structural organization. *In vitro* work already showed that phosphorylation of vimentin by cAMP-dependent protein kinase A soften the filaments (*44*). Moreover, it would be also relevant to determine whether other IF types, such as keratins or neurofilaments, also possess an axial repeat and also unfold their alpha-helices cooperatively during filament stretching. Recent *in vitro* work showed that keratin filaments have different mechanical properties than vimentin, and that keratin subunit might slide past each other upon stretching unlike vimentin (*45*). These results, added to fact that keratin filaments are made of ULF with only 4 tetramers per filament cross-section (*46*), suggest that keratin and vimentin filaments have different structural organizations. Because the vimentin rod domain is clearly shorter than that of keratins, we expect keratins to have longer axial repeats than vimentin, as also predicted by theoretical modeling (*43*). The difference between keratins and vimentin molecular organization could explain why the two types of filaments form disjoint networks that cannot copolymerize (*7, 43*). On the contrary, as vimentin, nestin and GFAP have been shown to co-polymerize at the level of single filament (*47, 48*) and to form heteropolymers (*49, 50*), we expect GFAP and nestin to display a similar axial repeat as vimentin.

IFs have the unique ability to reversibly elongate along their length (*14–17*). In particular, Block et al, used double optical trapping of individual purified filaments to probe their viscoelastic properties (*16*). They showed that although the filament stretching was reversible, vimentin filaments mechanical properties change after stretching, displaying stretch-softening (*16*). Their theoretical modeling combined with stochastic simulations suggested that the dependence of mechanical properties on deformation history may result from the unfolding of alpha helices in the rod domain. Our results provide further experimental evidence that it is indeed conformational changes which are responsible for the stretchability of vimentin IFs. By stretching isolated filaments attached to an elastic subtract via antibodies, we observed an increase of the axial repeat which corresponds to the distance between consecutive tails domains. However, the aspect ratio of the Spots, which corresponds to the intertwined tail domains, is the same without or with stretching, indicating that the tails remain together. These results suggest that the mechanical reorganization within ULF occurs within the rod domains, and not within the intertwined regions. Therefore, the molecular mechanism involving unfolding of alpha helices is more likely than sliding of individual vimentin tetramers, which would lead to a disordering/dislocation of ULFs and degradation of the axial repeat. Moreover, this model is compatible with the reversibility of filament elongation with force (*14–17*). However, live SRM with repeated cycles of strain will be required to further investigate the molecular mechanism responsible for the tensile memory of vimentin IFs.

Although the vimentin network is known to protect cells from large deformations (*5*), we found that vimentin filaments were not extended inside MEF cells for up to 50% amplitude of cell stretching. Instead, filaments reoriented along the stretch axis, with a mechanism which may implicate interaction with other cytoskeletal elements (*47*). This lack of elongation is intriguing because similar stretching amplitudes had a stronger impact on other structures, triggering disassembly of caveolae (*51*), or ruptures in actin stress fibers (*29*). Furthermore, 4-10 % stretch was sufficient to trigger mechano-sensing and mechano-transduction in integrin-based adhesion and to trigger active remodeling of the actin cytoskeleton that deform and recruit proteins in mechanosensitive structures (*29*). One possible explanation is that vimentin filaments are not under tension before stretching. Indeed, the requirement to stretch filaments is that longitudinal forces are exerted on at least two positions and that the filaments are already under tension. If the force is applied to only one position, for example via connection to integrin-based adhesions (*31*), they will not be extended. If the force is applied to more than one position but the filaments are folded or fluctuate between the positions where the force is applied, stretching will first lead to pulling out the slack in the filaments before extension along their length. However, although we did not observe an axial repeat increase on average, we cannot exclude that a small fraction of the filaments was stretched locally. In this case, this local stretching would have a limited contribution to the overall pairwise distances parallel to the stretch axis. The other explanation is that IFs mechanical connections within the cells are slipping, quickly releasing the tension during the stretch. Thus, our results illustrate the fact that vimentin filaments participate in mechanical energy dissipation during 50% cell stretching, behaving as stress absorbers (*5*). These mechanical properties may also be relevant to other types of IFs such as desmin, another type III IF protein found in highly deformable muscle cells (*7*), or keratins that protect epithelial cells from damage or stress (*4*). More work will be also necessary to understand how the force applied during cell stretching is propagated to cytoplasmic IFs.

In conclusion, the description of the vimentin structural organization by SRM is a powerful tool to investigate the molecular mechanisms involved in filament assembly and response to mechanical strain. In the future, it will be interesting to investigate quantitatively to what extent the size of the axial repeat scales with the tension applied on the filaments and to explore in more detail the coordination between the different subunits within filaments, both *in vitro* and in cells. Since cells are subjected to complex deformation history, it will be also important to apply more complex stretching patterns with controlled amplitudes, frequencies and durations. Moreover, testing if the axial repeat is present in other types of IFs and if it could be impacted by post-translational modifications (*18*) or IF protein mutations related to diseases (*52*) would be of prime interest. We anticipate that our approach will open the door to new studies aiming at elucidating the role of IFs in cell mechano-sensing and more generally in cell adaptation to external stimuli.

## Materials and Methods

### Reagents

GFP-nanobodies labeled by Alexa Fluor 647 (GFP-nanobodies-AF647) were purchased from Chromotek (# gb2AF647, Planegg-Martinsried, Germany) and Nanotag (FluoTag-X4 # N0304). Spot-nanobodies (# etb-250) and Star635P GFP-nanobodies (# gbas635p) were purchased from Chromotek. Anti-vimentin antibody, monoclonal, clone V9 was purchased from Sigma-Aldrich (Cat. # V6389, St. Louis, MO, USA). Anti-Mouse antibody AF647 was purchased from Jackson Immuno Research. All the chemicals were purchased from Sigma-Aldrich, except specifically noted.

### Labeling of Spot-nanobodies with AF647 or AF594

Spot-nanobodies were labeled by Alexa Fluor 647 for dSTORM or Alexa Fluor 594 for STED using a NHS ester labeling kit from Molecular Probes (#A37573 and # A37572 respectively) following the protocol provided by the company. The buffer of the nanobodies was changed to 100 mM sodium bicarbonate pH8.3 using zeba columns (ThermoFisher, Waltham, MA USA). The amine-reactive dye was dissolved in DMSO at 10 mg/mL just before the use, and diluted 10x in the nanobodies solution. After 1h incubation and stirring, the excess of dye was removed using the dye removal column from Pierce (#22858, ThermoFisher).

### Vimentin constructs

Mouse Vimentin-GFP (pEGFP-N3-Vimentin) plasmid was kindly given by Danielle Pham-Dinh and described previously (*53*). The vimentin constructs (Y117L mutants and addition of a Spot tag) were produced using a Q5 Site-Directed Mutagenesis kit (Biolabs cat # E0554S) using the primer lists in Table 1. Spot-vimentin-GFP was obtained by addition of a Spot tag (sequence PDRVRAVSHWSS (*37*)) at the N-terminal of pEGFP-N3-Vimentin. Spot-Vimentin was obtained by chopping the GFP at the C-terminal end of the pEGFP-N3-Vimentin construct and then adding a Spot tag between the rod and tail domains using linkers made of 2 amino acids (SG). Vimentin-Spot plasmid for bacterial expression was cloned from the human vimentin WT plasmid kindly provided by Harald Herrmann (*11*). The cloning procedures followed strictly the protocol provided by Biolabs (PCR with the initial construct and the primers written in the table listed below, ligation for 5 min at room temperature, bacteria transformation). About 4 clones were selected per condition and sequenced. The Vimentin-HALO construct was obtained after cloning using the HiFi DNA Assembly Cloning Kit (Biolabs cat # E5520S) starting from the tubulin-HALO plasmid (addgene #64691) for the vector and the pEGFP-N3-vimentin-GFP construct mentioned above for the insert. The cloning procedure followed strictly the protocol provided by Biolabs (PCR of the vimentin insert, PCR of the HALO vector, ligation with twice more insert than vectors, and then transformation). The primers used are listed in Table 1.

**Table 1:**
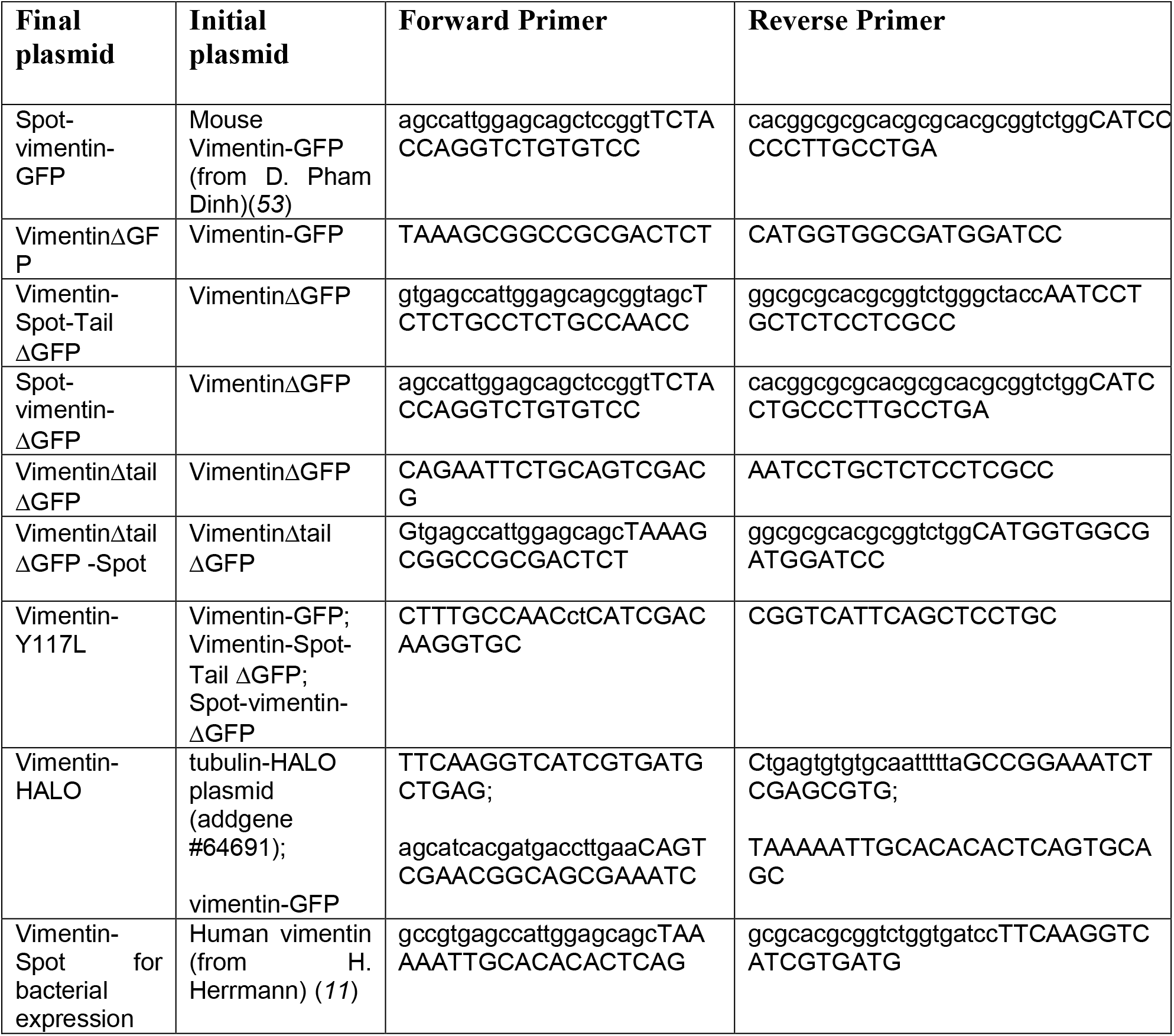
List of primers from cloning the vimentin constructs

### Cell culture, nucleofection

MEF cells vimentin KO described in (*30*) (kind gift from Harald Herrmann, DKFZ, Germany) and the CRISPR cell line U2OS Nup96 – SNAP (kind gift from Jonas Ries, EMBL Heidelberg, Germany) were grown in Dulbecco’s modified Eagle’s medium (DMEM) with 4.5 g/L glucose and supplemented with 10% FBS, 1% penicillin-streptomycin (pen-strep) and 1% non-essential amino acid (NEAA) at 37 °C in 5% CO2. For MEF vimentin KO cells, starting from a 10 cm diameter petri dish, cells grown to confluence were trypsinized and electroporated with a Nucleofector machine (Lonza). We used 2 µg of DNA, mixed with 97 µL of solution for electroporation (Mirus MIR 50111). After electroporation, cells were resuspended in pre-heated medium and plated on clean 18-mm n°1.5H glass coverslips (Marienfeld) in 12 wells plate with ~3000 cells per well. U373 cells, which are astrocytoma cells, were grown Minimum Essential Medium (MEM) supplemented with 10% FBS, 1% penicillin–streptomycin and 1% non-essential amino acids. DMEM, MEM, Pen-strep, NEAA and FBS were purchased from Gibco.

### Drug treatments

Nocodazole was added at a final concentration of 10 µM, 1h before fixation (stock at 10 mM in DMSO). After 1h incubation with nocodazole, vimentin filaments did not have the time to collapse around the nucleus (*47*). Blebbistatin was used at a final concentration of 2 µM (stock at 10 mM in DMSO), and was also added 1h before fixation. Nocodazole was purchased from Calbiochem (San Diego, CA, USA), and Blebbistatin from Sigma-Aldrich.

### Fixation and labeling

24h after electroporation, cells were fixed in cold (−20°) methanol for 5 minutes and blocked with 5% BSA in PBS for 1h. Cells were then incubated for 2 hours with fluorescent nanobodies (dilution 1/200e in PBS) at room temperature and rinsed 3 times with PBS for 5 minutes on an orbital shaker. For immunofluorescence, cells were incubated for 1 hour with anti-vimentin antibodies (dilution 1/500e in PBS) after blocking, washed three times 5 minutes in PBS and incubated for another hour with anti-mouse AF647 antibodies (dilution 1/500e in PBS). After 3 washes of 5 minutes in PBS, cells are incubated with 100 nm tetraspeck beads (#T7279, ThermoFischer) with a dilution of 1/500 in PBS for 30 minutes on an orbital shaker at room temperature and then stored at 4° in PBS before use.

### Sample preparation, Imaging buffer

18 mm coverslips are mounted on magnetic sample holders (Live Cell Instrument, CM-B18-1), with 1 mL of imaging buffer, and then closed with a lit. The imaging buffer is made of Tris-NaCl buffer (50 mM Tris, 10 mM NaCl, pH = 8) supplemented with 10% of glucose (100 mg/mL), 10 mM MEA, 0.5 mg/mL (75 U/mL) glucose oxidase and 40 μg/mL catalase (80 - 200U/mL). For dSTORM images of vimentin filaments reconstituted *in vitro*, the concentration of NaCl was increased to 100 mM.

### 2D-dSTORM imaging and processing

2D-dSTORM images were acquired on a Elyra Zeiss microscope, using the TIRF µ-HD mode which focus the intensity of the lasers in the center of the field of view, a x100 NA 1.46 objective and a 1.6 optovar lens and a 512×512 EMCCD camera. We worked with 20×20 µm^2^ field of view, used 15-20 ms exposure time and acquired 20 to 40 000 frames at maximal intensity of the 642-nm laser power to pump most of the Alexa Fluor 647 dyes to the dark state (~2 kW/cm2) as described previously (*54*). Super-resolution images were reconstructed and drift correction was carried out using the Picasso software available on GitHub and previously described (*24*).

### 3D-dSTORM imaging

We used a home-made microscopy system to perform 3D-dSTORM imaging, based on a previously published method (*38*). The system consists of two microscopes facing each other, oriented horizontally and organized around the biological sample. The first microscope is composed of (i) a high numerical aperture objective lens (Olympus - UPSALO100x - NA1.4), (ii) a Semrock-Di01-R405/488/561/635-25×36) quadband dichroic that reflects the excitation laser beam coming from a near infra-red laser light (λ=642nm Oxxius LBX series) and filters the fluorescence light coming from the sample, (iii) a tube lens composed of an achromatic doublet (f=250mm Thorlabs-AC508-250-A-ML) and a 4f system composed of two achromatic doublet lenses (f=100mm-

Thorlabs AC254-100-A-ML) used to image the biological sample on the ship of an EMCCD camera (CAM1-512×512px Andor Ixon 897). In the intermediate plane of the 4f system, a cylindrical lens (f=1m Thorlabs - LJ1516RM-A) is added, modifying the shape of the point spread function. A long pass and a band pass filters Semrock (Long wave Pass 647 and FF01-676/29) are added in front of the camera to filter out residual photons from the laser. The second microscope is identical to the first one, balancing the number of photons collected.

The biological sample is sandwiched between two high resolution coverslips (#1.5H, 224×50 mm, Marienfield) which are placed between the two objective lenses and held on a home-made 3D sample holder. Fluorescent beads are added to the biological sample in order to help select the same focal plane with both microscopes and to correct *a posteriori* the 3D spatial drift.

Once the focal plane is selected in the biological sample with the both microscopes, several tens of thousands of 60×60 µm images are acquired by the two synchronized cameras. Due to the geometry of the system, the astigmatic PSFs are oriented at 90° from each other on both cameras. Single molecules appearing in successive images are localized with ZOLA-3D (*55*). This algorithm allows the correction of 3D spatial drift using fiducial markers and has been adapted to this imaging system. Optionally, ZOLA-3D can correct for sub-pixel displacements between the images acquired by the two cameras and a specific model has been developed to create a localization table combining the astigmatic PSFs acquired in both cameras ensuring the best localization accuracy.

### STED imaging

STED images were acquired on a STEDYCON microscope, and an expert line microscope from Abberior Instruments GmbH, using the adaptive illumination technology reducing laser illumination of the sample (around 5-10x). Spot-vimentin Y117L-GFP proteins were labeled with GFP-nanobodies Star 635P and Spot-nanobodies Alexa Fluor 594, following the protocol described above.

### In vitro reconstitution of vimentin-Spot filaments

Vimentin-Spot protein was purified from bacteria as described previously (*40*). Briefly, vimentin-Spot was expressed in BL21 star (Sigma-Aldrich) in Terrific Broth medium overnight at 37°, after induction at a DO of 1.2. After centrifugation, bacteria were lysed with lysozyme in the presence of DNase (Roche), RNase (Roche) and protein inhibitors (pefabloc and PMSF). Inclusion bodies were collected and washed 5 times by successive steps of centrifugation and resuspension using a cooled douncer. After the last washing step, inclusion bodies were resuspended in a denaturing buffer (8M urea, 5 mM Tris pH 7.5, 1 mM EDTA, 1 mM DTT, 1% PMSF) and centrifuged at high speed (100 000 g) for 1h. Starting from the supernatant, Vimentin-Spot purification was obtained after two steps of exchange chromatography, using first an anionic (DEAE Sepharose, GE Healthcare) then a cationic (CM Sepharose, GE Healthcare) column. The protein was collected in 2 mL tubes, and the concentration was monitored by Bradford. The most concentrated fractions were pooled together and stored at −80° with additional 10 mM methylamine hydrochloride solution. Vimentin-Spot proteins were renatured after stepwise dialysis (8M, 6M, 4M, 2M, 1M, 0M urea) into sodium phosphate buffer (2.5 mM, pH 7) supplemented with 1 mM DTT, using dialysis tubing (Servapor, cut off at 12 kDa). Each step lasted 15 to 30 minutes, except the last dialysis performed overnight at 4° with 2 L of buffer. Filament assembly was triggered by addition of 100 mM KCl (final) to a solution of renaturated vimentin-Spot at 0.2 mg/mL, and elongation was performed by incubation at 37° for 3h for long filaments and 2 seconds to obtain ULFs.

### Transmission electron microscopy imaging of reconstituted filaments

Vimentin-Spot filaments were grown at 0.2 mg/mL for 3h at 37° after addition of 100 mM KCl and then fixed with an equal volume of glutaraldehyde 0.5% in a sodium phosphate buffer (2.5mM) supplemented with 100 mM KCl. 4 microliters of the sample were Spotted on a carbon coated grid primarily glow discharged and incubated at room temperature for one minute. 2% uranyl acetate in water was used to contrast the grids and incubated for one minute. The grid was then dried and observed under 120kV using a Tecnai microscope (Thermofisher) and imaged using a 4kx4k Eagle camera (Thermofisher).

### Stretching experiments coupled to DNA-PAINT, imaging vimentin-HALO in cells

The stretching device and its coupling to DNA PAINT was previously described (*29*). Briefly, a plasma-cleaned PDMS sheet (10 µm; Sylgard 184; Samaro DE9330) was deposited on a plasma-cleaned glass coverslip, lubricated by a thin layer of low-viscous glycerol, and reinforced by a thicker PDMS elastomer frame. The uniaxial stretch was applied thanks to a 3D printed device consisting of a fixed holding arm and a mobile arm connected to a piezo-electric motor (M-663 Linear Positioning Stage; PI). PDMS substrate was coated with human fibronectin for 90 minutes at 37°. Then, 18h after electroporation with a mix of vimentin-HALO:vimentin-GFP 9:1, MEF vimentin-KO cells were plated on the stretching device and spread for 3h at 37°C. The device was mounted onto the microscope at 37°C in order to acquire low resolution vimentin-GFP images of cells before stretching. Then, at room temperature and outside the microscope, a single uniaxial stretch of 30% or 50% was applied with a speed of 0.5 mm/s and a duration of 5 or 10 seconds (respectively), followed by rapid cell fixation in 4% paraformaldehyde, 0.3% Triton and 0.3% glutaraldehyde in PBS buffer for 20 min at 37°. Simultaneously, the stretching arm was clamped to the fixed arm using a thread and groove system, in order to maintain the stretching throughout the subsequent labelling and imaging steps. Then, cells were then blocked for 90 minutes with 3% BSA in PBS, and incubated with the HALO-Paint probe for 15 minutes. Back on the microscope, we located the cell imaged before stretching and acquired low resolution images of vimentin-GFP after stretching. Then super-resolution imaging was performed. Cells were imaged at 25 °C in the stretching device with an inverted motorized microscope (Nikon Ti) equipped with a CFI Apochromat TIRF 100× oil, NA 1.49 objective and a perfect focus system (PFS-2), allowing long acquisition in TIRF illumination mode. For DNA-PAINT microscopy, cells expressing Vimentin– HALO were first incubated with 90 nm gold nanoparticles (Cytodiagnostics) to serve as fiducial markers. Vimentin-HALO was visualized with Cy3B-labelled DNA imager strands added to the stretching chamber at variable concentrations (2-5 nM) using a streaming mode at 6.7 Hz. Cy3B-labelled strands were visualized with a 561-nm laser (Cobolt Jive). Fluorescence was collected by the combination of a dichroic filter and emission filters (dichroic: Di01-R561; emission: FF01-617/73; Semrock) and a sensitive scientific complementary metal–oxide–semiconductor (ORCA-Flash4.0; Hammamatsu). The acquisition was steered by MetaMorph software (Molecular Devices) in streaming mode at 6.7 Hz. Vimentin–GFP was imaged using a conventional GFP filter cube (excitation: FF01-472/30; dichroic: FF-495Di02; emission: FF02-520/28; Semrock). DNA-PAINT image reconstruction and drift correction were carried out using the Picasso software (*24*).

### Stretching experiments coupled to 2D-dSTORM, imaging reconstituted vimentin-Spot filaments

Using the same stretching device and microscope setup described above, the thin plasma-cleaned PDMS layer was incubated with vimentin antibodies (Santa cruz (V9): sc-6260, conc : 0.2 mg/mL) dilution 1/10 in buffer V (2.5 mM sodium phosphate pH 7, 100 mM KCl) for 15 minutes, rinsed with buffer V, incubated with a mix of vimentin-Spot filaments diluted 50 times, tetraspeck beads diluted 500 times and Spot-nanobodies-AF647 diluted 1000x in buffer V for 3h. Then the stretching chamber was rinsed with STORM buffer and filled with 2 mL of STORM buffer freshly prepared and changed every 30 minutes. A single uniaxial stretch of 30% was applied with a speed of 0.03 mm/s and a total duration of 1 minute.

### ULF pairwise distance analysis

For 2D measurements: first the ULFs were detected in low resolution images by thresholding. Then for the isolated ULF dots which contained only two peaks in the super-resolution images, intensity profiles were plot along a thick line (5 pixel of 5 nm) containing the two peaks, and the distance the peaks, called pairwise distance, was quantified using a double Gaussian fit.

For 3D measurements: the beginning of the procedure is the same. ULFs are detected from low resolution images by thersholding. We analyzed the ULFs when two peaks were present within the ULF dots in the 2D projection of the 3D super-resolved stack. A transverse section was plot along the doublet (as shown in Fig. 1G side view). The pairwise distance was calculated as the distance between the centroids within the side view. Note that when ULFs are close to being vertical, the 2D projection of the pairwise distance goes below the resolution of technique (~ 25 nm) and are therefore not detected. The tilt angles of ULFs close to 90° are therefore underestimated with this method.

### Filament pairwise distance analysis

Analysis of the double length was performed using MATLAB (The Mathworks) and Fiji and involved several steps depicted in Fig. 3C: (1) isolating filaments, (2) detecting single vimentin peaks and liking them to their 2 nearest neighbors, (3) removing graph nodes that have 4 edges or more, (4) resolving triplets, (5) removing the connected components that have fewer than 4 nodes, and (6) measuring distances from the remaining edges.

#### 1) Isolating filaments

The reconstructed super-resolution image of vimentin contains spurious signals away from filaments that we need to filter out. Because true vimentin peaks on filaments are spaced by a distance smaller than the filament persistence length, we can use filter operations to yield a broad mask that contains only the filaments. To do so we used the *Tubeness* filter present in Fiji (*56, 57*), with a σ value larger than the peak separation, but smaller than the filament persistence length would work. We picked σ = 12 pixels. We then thresholded the tubeness image, and segmented it into individual regions. We rejected regions with an area smaller than 1000 pixels and with an aspect ratio larger than 0.5. The resulting mask delineates individual filaments. Some high-curvature filament locations were also filtered out by this method.

#### 2) Detecting single vimentin peaks and liking them to their 2 nearest neighbors

In the resulting image, a single vimentin peak is made of possibly several bright Spots, clustered around the vimentin peak location. We tried several approaches to yield the vimentin peaks location from individual Spots. We first tried using the DBSCAN algorithm on the emitter locations directly, as first proposed in Endesfelder et al., 2013 (*58*), and with an improved implementation proposed in Tran et al., 2013 (*59, 60*) to harness close clusters. However, in our case the number of individual Spots per vimentin peak appears too small to achieve satisfactory results. Individual clusters bridge over several vimentin peaks. We therefore turned to direct analysis of the reconstructed super-resolution image. We finally settled to harvest peak locations by directly detecting local maxima in the reconstructed image, filtering for the brightest ones and refining their location using a simple centroid calculation over 5×5 pixels. We then built a simple undirected graph connecting all vimentin peaks closer than 150 nm, and used it to further filter out spurious peaks.

#### 3) Removing graph nodes that have 4 edges or more

Ideally the graph would be made of long connected components corresponding to filaments, with each vimentin peak having 1 or 2 connections to the neighbor. However, filaments can cross and spurious detections complicate the analysis. We first filtered out all peaks that have 4 or more connections to neighbor peaks (quadruplets). Some peaks had 3 or more connections (triplets), which we tried to resolve in the following manner:

#### 4) Resolving triplets

A peak with 3 connections to 3 neighbor peaks corresponds to 3 candidate doublets. We look for the angles between each doublet pair. If the doublets are not roughly aligned, that is, their angle is below 130° and above −130°, they make a sharp nick. We then vote against these 2 doublets. We do this for all the possible doublet pairs (3 combinations are possible). And in the end, we look if there is a doublet that was voted against twice. This is the doublet that is not aligned with the roughly straight line built by the two others. We then remove it. If there is no such doublet (the 3 of them have 0 votes, because they are all roughly aligned), then we simply remove the longest one.

#### 5) Removing the connected components that have fewer than 4 nodes

The remaining peaks all have 1 or 2 connections to neighbor peaks. In the graph they build several connected components representing parts of individual filaments. Some components are just made of a few peaks, and for which defining a global orientation is hard. If such a component is made of less than 4 nodes, we remove it.

#### 6) Measuring distances from the remaining edges

The remaining edges in the graph correspond to single vimentin doublets, belonging to long filament parts, from which we extract the doublet length.

In a typical image used in this analysis (DNA PAINT as shown in figure 3), we find 38890 peaks in step ii) of which we keep only the 29524 brightest. From them we build a graph made of 29524 nodes and 28379 edges. By filtering out quadruplets we remove 413 peaks. By curating the triplets, we remove 5502 edges. By removing the filament parts smaller than 4 nodes we remove 10797 peaks. In the end we are left with 18314 peaks and 15422 doublets, from which we yield 15422 pairwise distance measurements.

### Statistical analysis

All experiments were repeated at least three times except specifically mentioned. The statistical analyses were performed using Graphpad. P values were calculated using a nested t-test of the three repeats for figure 1 D-J and 2C and a paired t-test for figure 3E. A *P*-value below 0.05 was considered as statistically significant * *P* < 0.05, ** *P* < 0.01, *** *P* < 0.001, **** *P* < 0.0001.

## H2: Supplementary Materials

Fig. S1: STED images of vimentin ULFs in cells

Fig. S2: Other examples of vimentin ULFs

Fig. S3 Localization of vimentin ULFs and vimentin filaments in MEF vimentin-KO cells.

Fig. S4: Nocodazole but not blebbistatin impacts vimentin ULF projected length probed by 2D-dSTORM

Fig. S5: Nuclear pores probed by 3D-dSTORM using a dual-objective microscopy setup with a cylindrical lens.

Fig. S6: Pairwise distances are not correlated with the mean amplitude of the doublet

Fig. S7: 3D-dSTORM image of vimentin filament within cells

Fig. S8: Microtubule depolymerization, fixation methods and cell types do not impact the distribution of pairwise distances between vimentin tails.

Fig. S9: Vimentin filaments exhibit a 49 nm axial repeat in the absence of vimentin tail and in endogenous network within astrocytoma cells

Fig. S10: Two-angle histograms of stretched cells with an amplitude of 33%

**Fig. S1:**
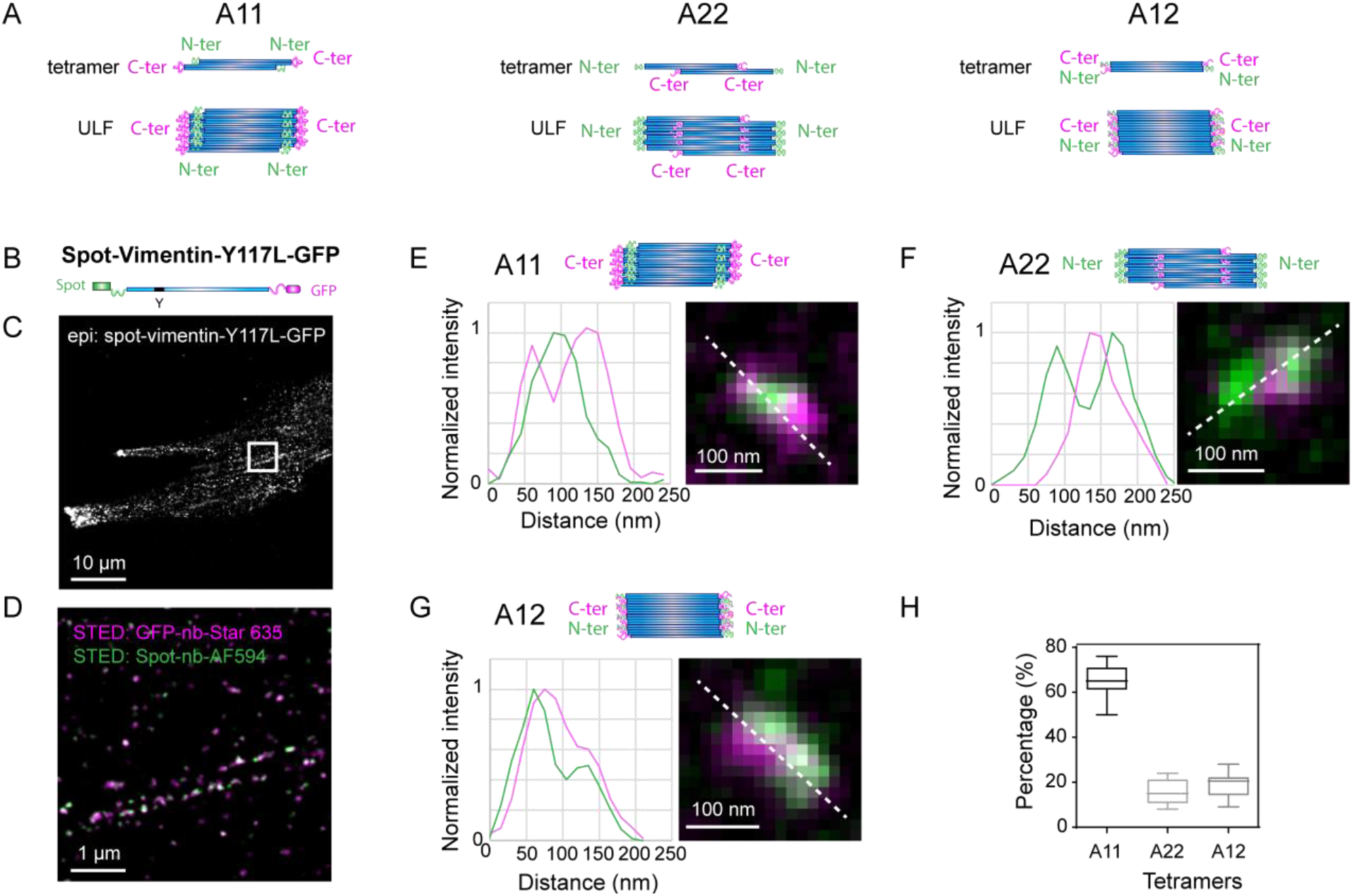
STED images of vimentin ULFs in cells. **(A)** Schematics of the different types of ULFs made of: A11 tetramers with the N-terminal ends inside and the C-terminal ends outside, A22 tetramers with the N-terminal ends outside and the C-terminal ends inside, and A12 tetramers with fully antiparallel dimers. **(B)** Schematic of the Spot-vimentin-Y117L-GFP construct with a Spot-tag at the N-terminal end and a GFP at the C-terminal end. **(C)** GFP epi-fluorescence image of a MEF vimentin-KO cell expressing the Spot-vimentin-Y117L-GFP construct. **(D)** STED image of the boxed area in (C). The cell is immune-labeled with GFP-nanobodies-STAR 635 (magenta) and Spot-nanobodies-AF594 (green). **(E-G)** Schematics of the different types of ULFs made of A11 (D), A22 (E) and A12 (F) tetramers. Examples of STED images of these different types of ULFs, with the corresponding intensity profiles in magenta for the C-terminal tails and green for the N-terminal heads **(H)** Box plot of the distribution of ULF types which could be determined: A11, A22, A12 tetramers. Boxes are calculated from about 400 ULFs imaged in 8 cells from 2 independent experiments.

**Fig. S2:**
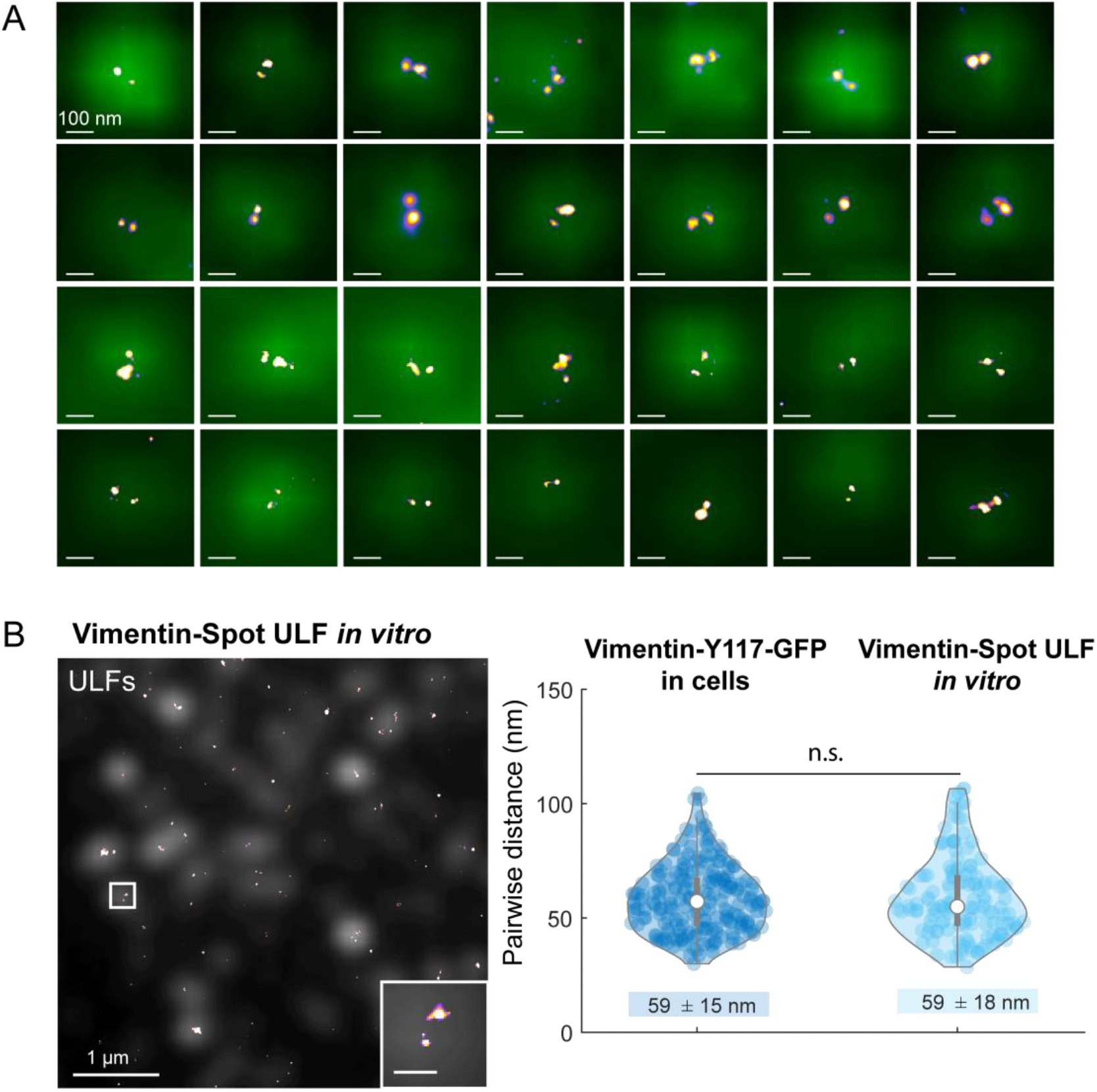
Other examples of vimentin ULFs in cells and *in vitro*. **(A)** Epi-fluorescence images overlaid with 2D-dSTORM images of GFP-nanobodies AF647. Examples of vimentin ULFs are extracted from 3 different cells. Vimentin-Y117L-GFP was expressed in MEF vimentin KO cells, which were fixed by cold methanol and immune-labeled with GFP-nanobodies AF647. **(B)** Left. Epi-fluorescence images (grey) superimposed on the corresponding 2D-dSTORM images (fire) of purified vimentin-Spot ULFs fixed 2 seconds after assembly started using glutaraldehyde. Right. Distributions of pairwise distances between vimentin-Y11L-GFP C-terminal measured in 3D (data from Fig. 2D “3D Control”). Mean values ± standard deviations of the total distributions are written below the violin plots. 204 and 141 ULFs were analyzed from 4 control cells acquired during 2 independent experiments and 3 independent *in vitro* experiments, n.s.: non-significant.

**Fig. S3.**
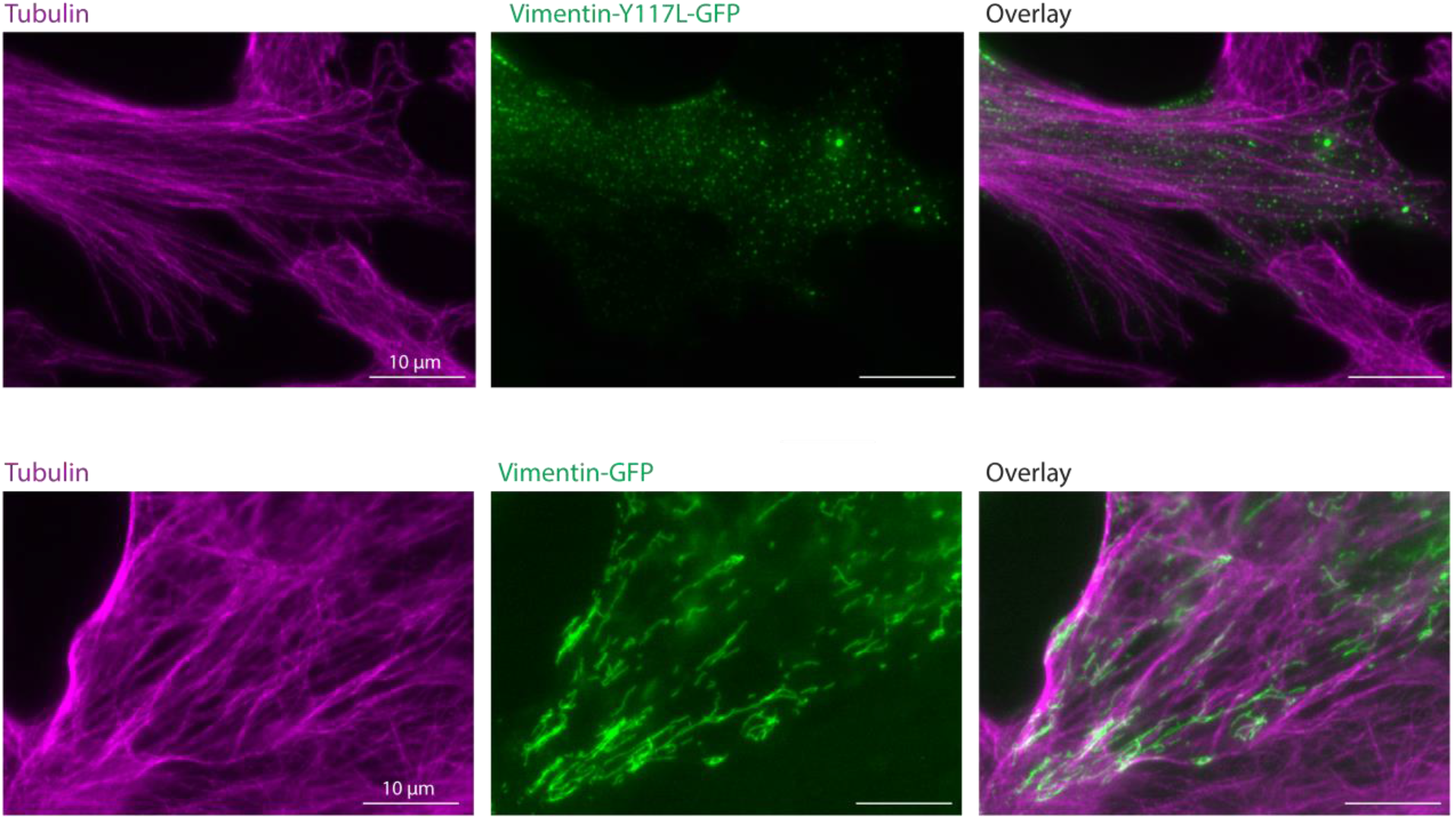
Localization of vimentin ULFs and vimentin filaments in MEF vimentin-KO cells. **(A)** Epifluorescence images of tubulin (magenta) and vimentin-Y117L-GFP (green) in MEF vimentin-KO cells fixed 1 day after nucleofection. **(B)** Epifluorescence images of tubulin (magenta) and vimentin-GFP (green) in MEF vimentin-KO cells fixed 1 day after nucleofection.

**Fig. S4:**
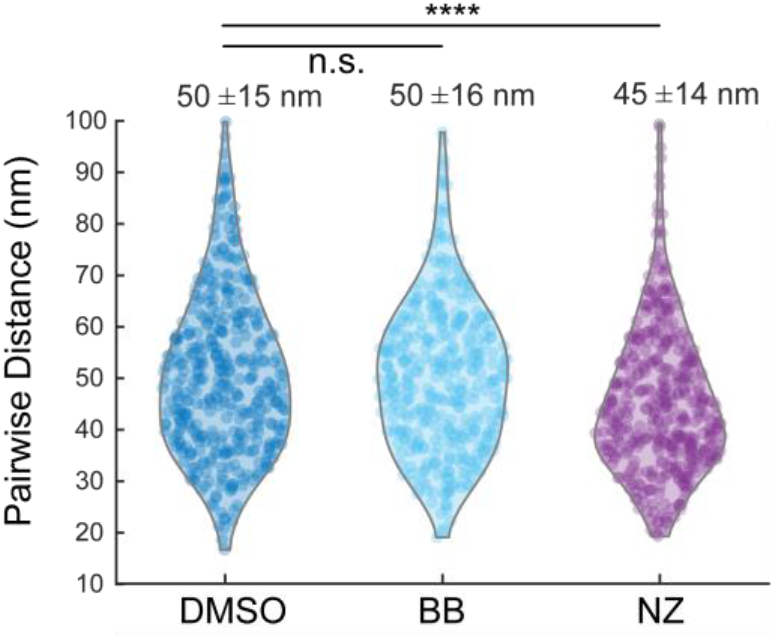
Nocodazole but not blebbistatin impacts vimentin ULFs projected length probed by 2D-dSTORM. Distribution of pairwise distances between vimentin-Y117L-GFP C-terminal ends labeled with GFP-nanobodies-AF647 in vimentin ULFs. Data were acquired by 2D-dSTORM in MEF vimentin-KO cells treated with DMSO (blue), blebbistatin (BB, cyan) and nocodazole (NZ, purple). Mean values +/- standard deviations of the total distributions are written above the violin plots. Data represent 518, 521 and 544 ULFs respectively, collected from 12, 15 and 14 cells from 3 independent experiments. P value was calculated with a nested t-test of the 3 repeats, ****: P<0.0001, n.s.: non-significant

**Fig. S5:**
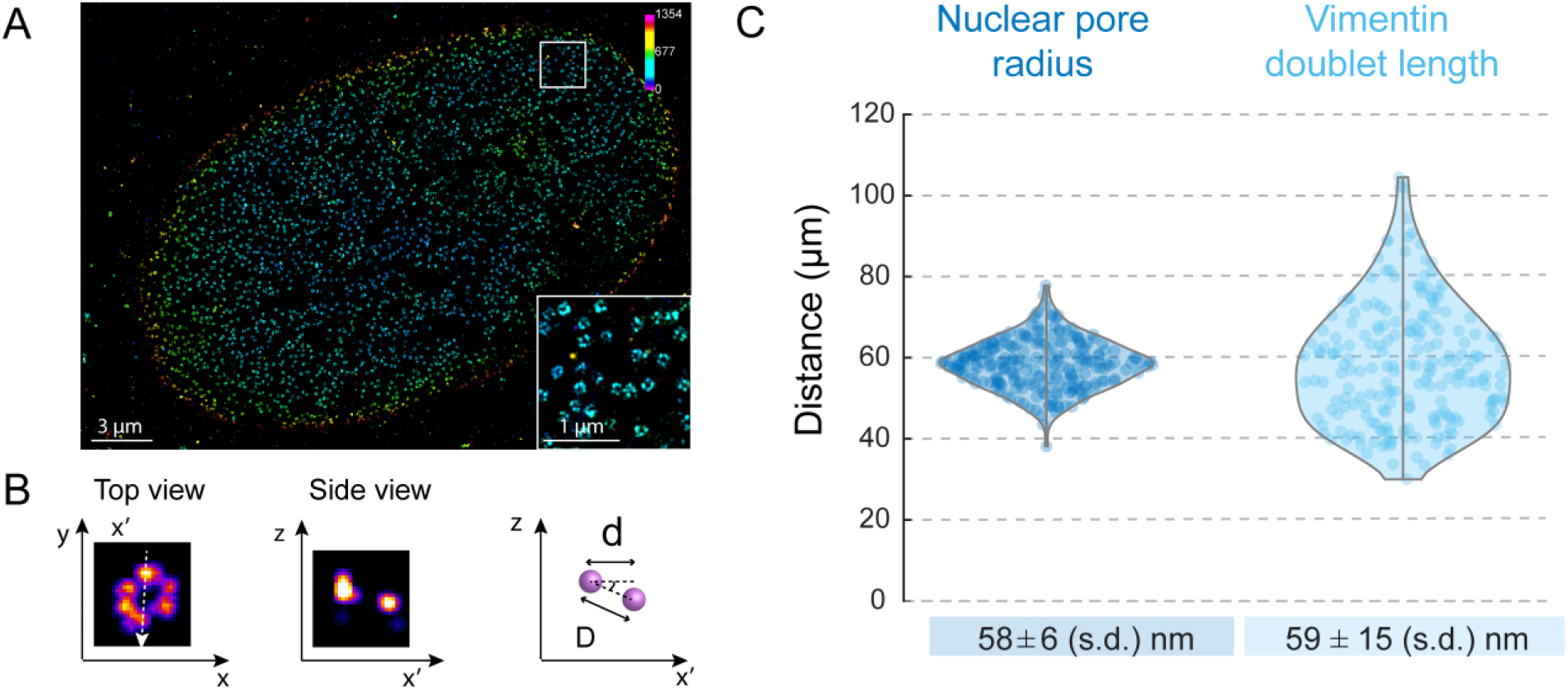
Nuclear pores probed by 3D-dSTORM using a dual-objective microscopy setup with a cylindrical lens. **(A)** 3D-dSTORM image of Nup96-snap labeled with SNAP-AF647, and expressed in U2OS cells as described previously (*34*). **(B)** Top and side views of a nuclear pore imaged by 3D-dSTORM, and a cartoon showing the projected diameter “d”, the real diameter measured in 3D, and the tilt angle. **(C)** Distribution of nuclear pore radius measured in 3D compared to vimentin doublet length (data from Fig. 2D, “3D control”). For nuclear pores, 231 measurements were performed from 2 independent cells. For vimentin doublets, 204 measurements were performed from 4 cells and 2 independent experiments. Data for nuclear pores and vimentin doublets were acquired on the same experimental setup and analyzed with the same method.

**Fig. S6:**
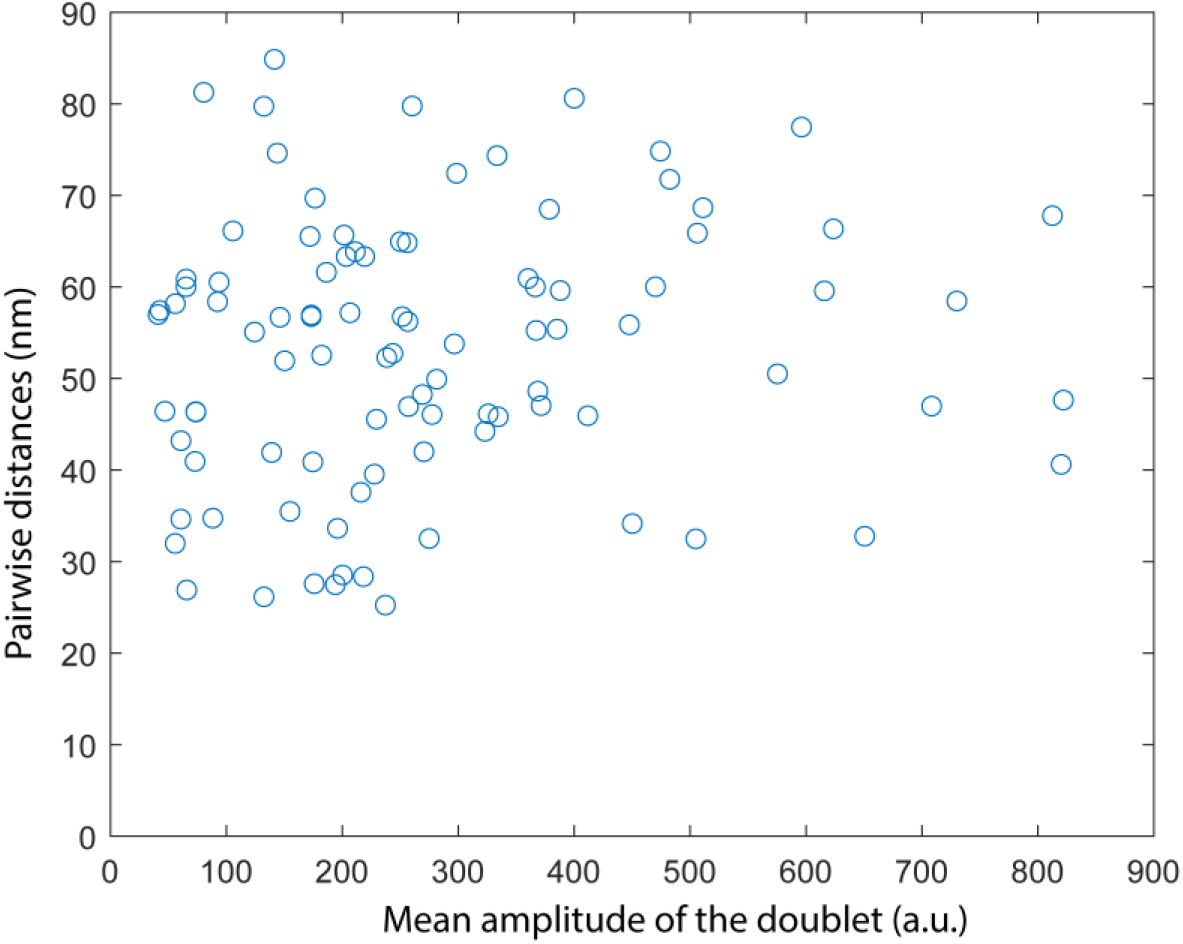
Pairwise distances are not correlated with the mean amplitude of the doublet. For each ULF observed in 2D-dSTORM after immune-labeling of vimentin-Y117L-GFP with GFP-nanobodies-AF647, we measured the pairwise distance using a double Gaussian fit of the intensity peaks and calculated the mean amplitude of the two intensity peaks. The Pearson correlation coefficient is 0.1 and the P value of the statistical test of no correlation against the alternative of a non-zero correlation is 0.3.

**Fig. S7:**
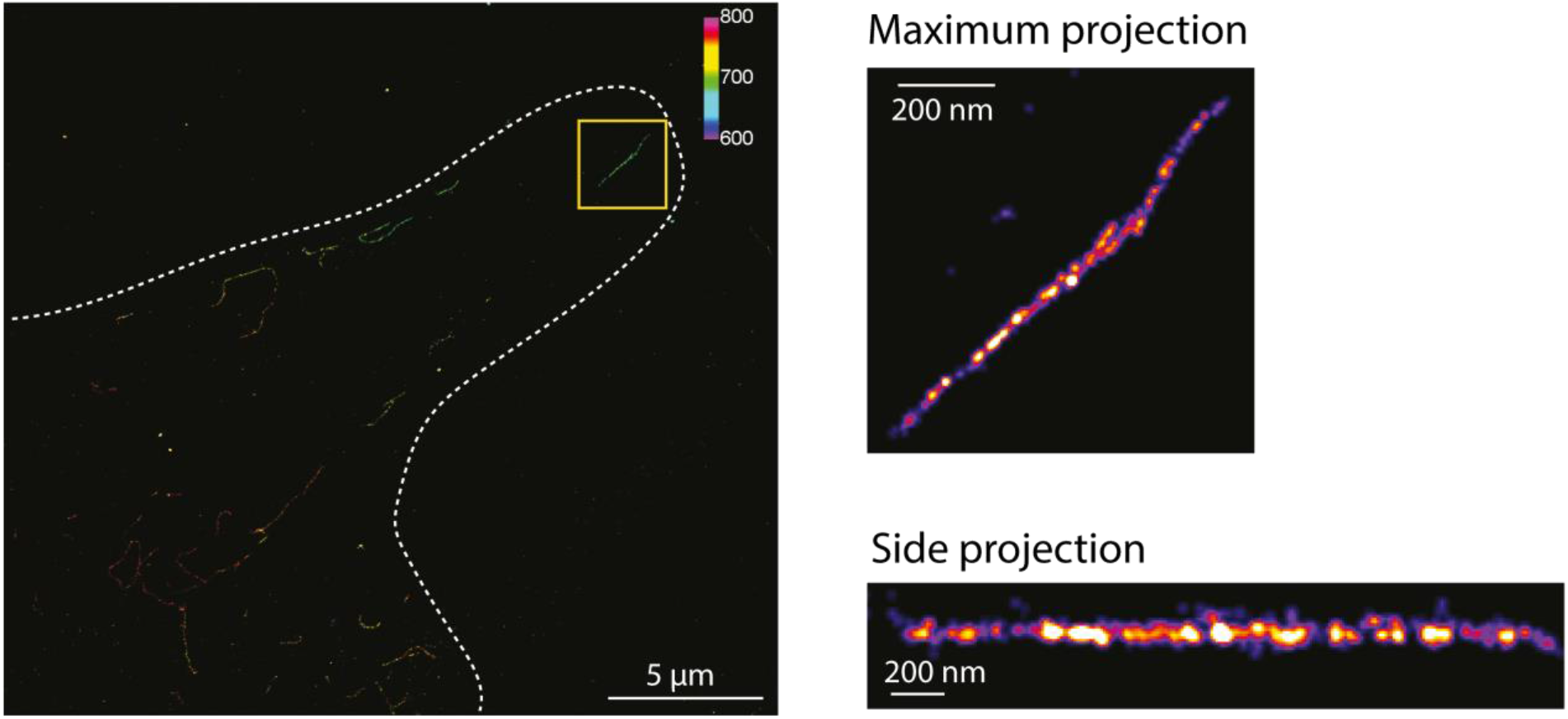
3D-dSTORM image of vimentin filament within cells. 3D-dSTORM image of a MEF vimentin-KO cell expressing vimentin-GFP and immune-labeled with GFP-nanobodies-AF647. Color scale bar in nm. Magnifications of the boxed region: top and side projections along the filament.

**Fig. S8:**
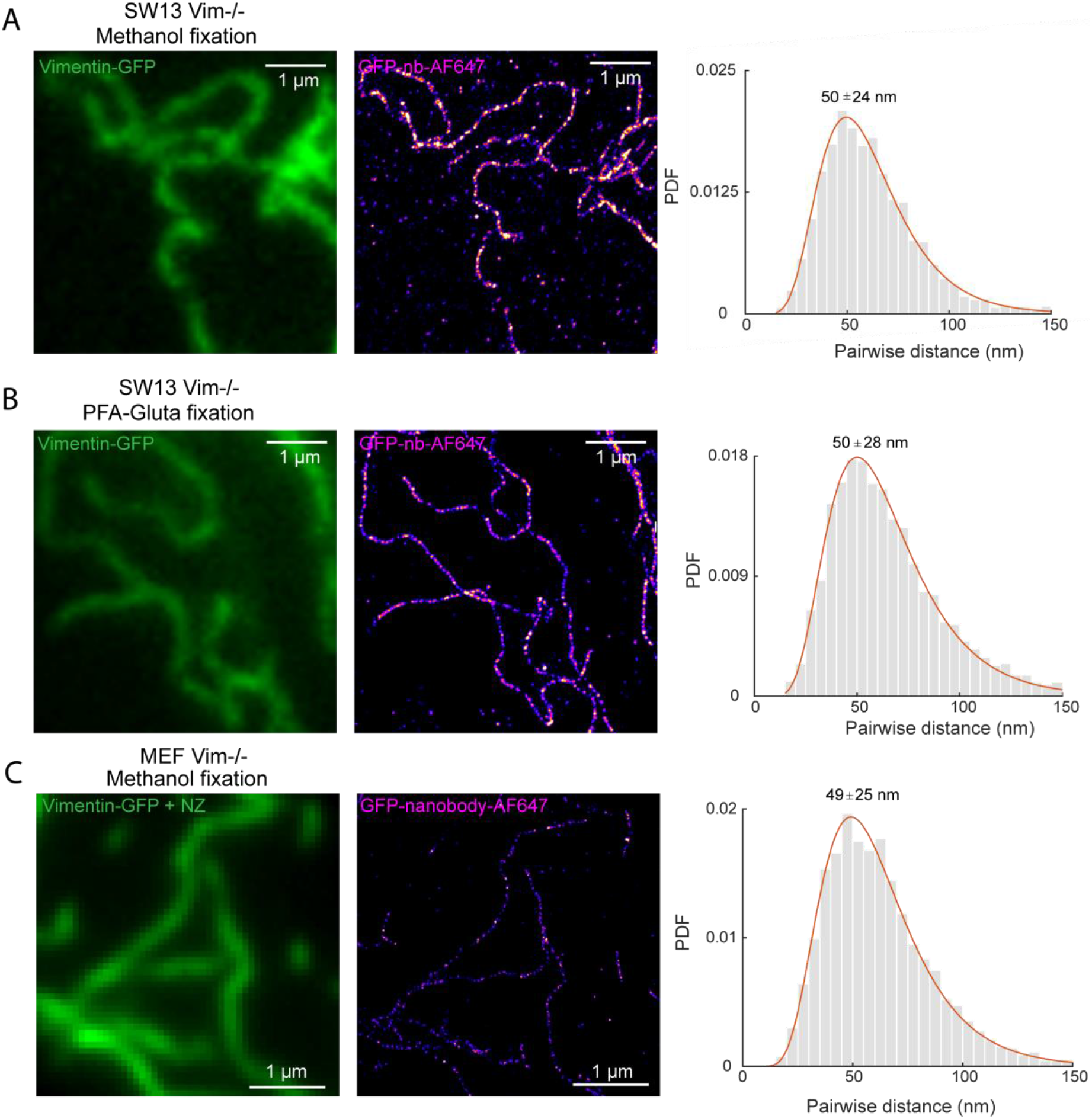
Microtubule depolymerization, fixation methods and cell types do not impact the distribution of pairwise distances between vimentin tails. **(A-B)** Left: epifluorescence image of SW13 vimentin −/− cells expressing vimentin-GFP and fixed with cold methanol (A) or a mix of PFA and Glutaraldehyde in PHEM buffer (B) (see methods). Middle: Corresponding 2D-dSTORM image of the GFP-nanobodies-AF647 attached to the GFP-C terminal ends of vimentin. Right: Histogram of pairwise distances detected automatically. Red: lognormal fit of the distribution. The distribution mode and standard deviation are written above the maximum. 2955 and 8999 pairwise distances were measured using data from 7 cells from 2 independent experiments for each condition respectively. **(C)** Left: Epifluorescence image of a MEF KO expressing vimentin-GFP and treated with 10 µM nocodazole before fixation with cold methanol. Middle: Corresponding 2D-dSTORM image of the GFP-nanobodies-AF647 attached to the GFPs. Right: Histogram of pairwise distances detected automatically. Red: lognormal fit of the distribution. The distribution mode and standard deviation are written above the maximum. 6398 pairwise distances were measured using data from 6 cells from 2 independent experiments.

**Fig. S9:**
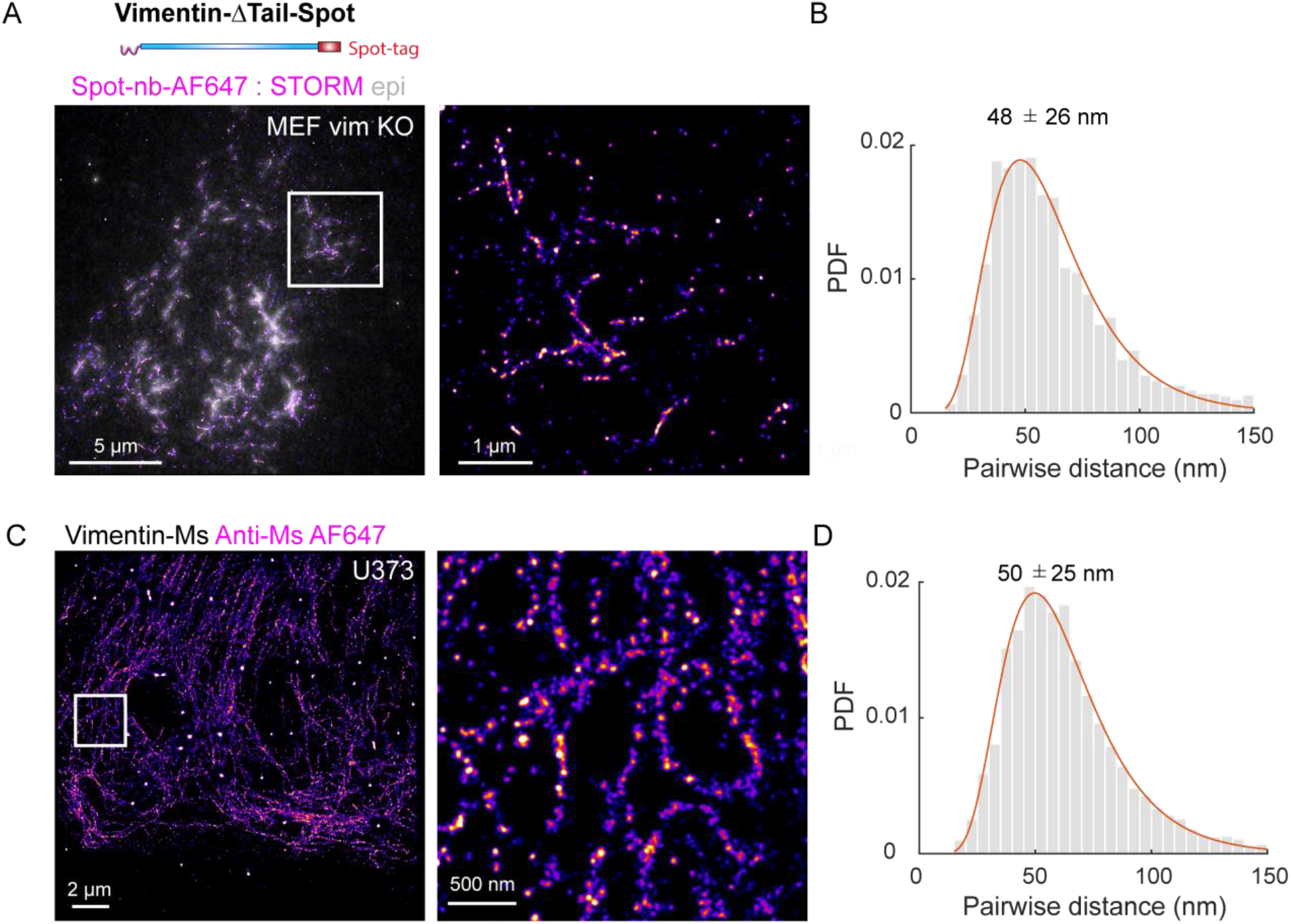
Vimentin filaments exhibit a 49 nm axial repeat in the absence of vimentin tail and in endogenous network within astrocytoma cells. **(A)** Top: Schematic of the vimentin-ΔTail-Spot construct. Bottom: epi-fluorescence and corresponding 2D-dSTORM images superimposed (fire) of a MEF vimentin-KO cell expressing the vimentin-ΔTail-Spot and immune-labeled with Spot-nanobodies-AF647. Magnification of the boxed region in the dSTORM image. **(B)** Histogram of the pairwise distances measured in dSTORM images of MEF vimentin-KO cells expressing the vimentin-ΔTail-Spot. The distribution mode and standard deviation are written above the maximum. 2207 pairwise distances were analyzed from 4 cells from 2 independent experiments. **(C)** 2D-dSTORM image of an astrocytoma U373 immuno-stained with anti-vimentin mouse primary antibodies (clone V9) and secondary antibodies labeled with AF647. Inset: magnification of the boxed region. **(D)** Distribution of pairwise distances measured automatically. Red curve corresponds to a lognormal fit. The distribution mode and standard deviation are written above the maximum. 10707 pairwise distances were analyzed from 8 cells from 3 independent experiments

**Fig. S10:**
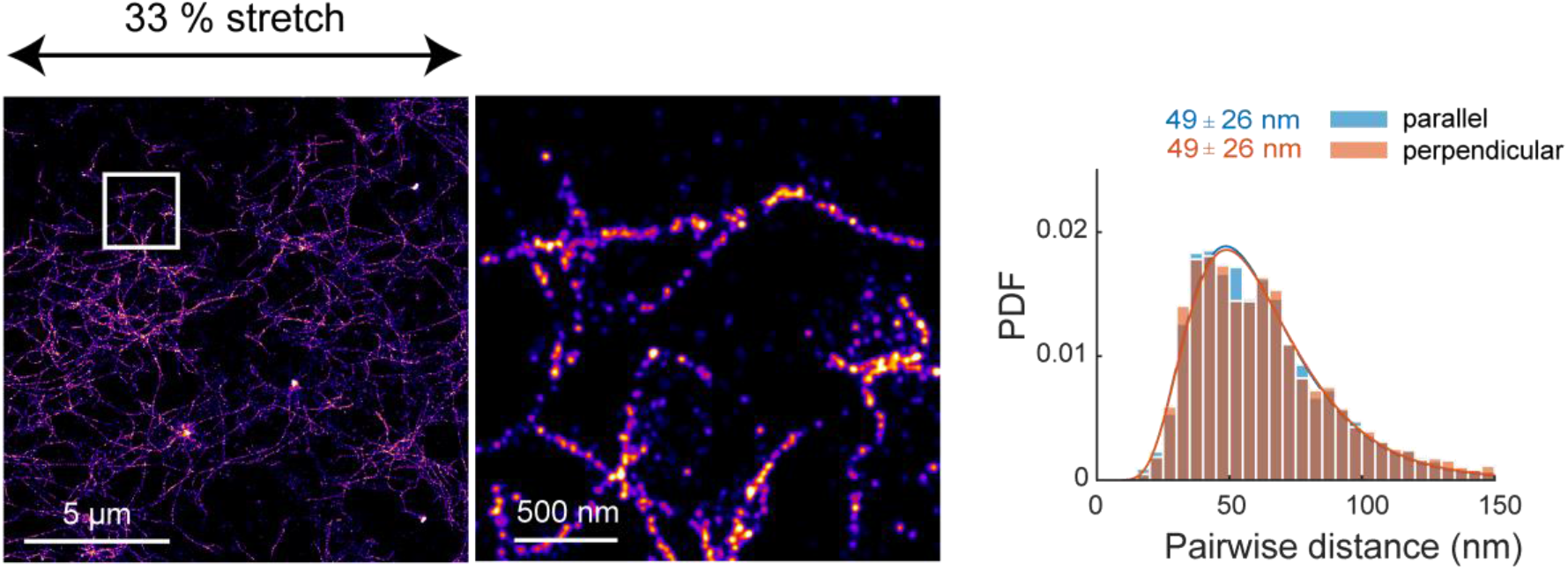
Two-angle histograms of stretched cells with an amplitude of 33%. Histograms of pairwise distances for pairwise distances aligned with uniaxial stretch (parallel, blue) and perpendicular to the stretch direction (perpendicular, orange), detected from the DNA-PAINT of vimentin-HALO with 33% stretch. Blue and orange curves correspond to a lognormal fit. The distribution modes and standard deviations are written above the curves. 4527 and 6017 pairwise distances were analyzed for perpendicular and horizontal distances respectively from 2 independent experiments.

## References and Notes

## Acknowledgments

We thank Rudolf Leube and Reinhard Windoffer from the university of Aachen, Pierre Nassoy from university of Bordeaux, Martin Lenz from university of Orsay well as Batiste Boëda, Vanessa Roca, Benoit Lelandais, Christian Weber and Thomas Obadia from Institut Pasteur for fruitful discussions, and Danielle Pham-Dinh, Jonas Ries and Harald Herrmann for reagents. The U2OS Nup96 – SNAP cell line used in Fig.S5 can be provided by Jonas Ries pending scientific review and a completed material transfer agreement. We thank Frédéric Eghiaian, Janina Hanne and Jan-Gero Schloetel from Abberior Instruments GmbH, for imaging our samples. Cécile Leduc thanks Harald Herrmann and his group for teaching her the techniques involved in vimentin purification, and the support of the EU-supported Cooperation in Science and Technology (COST) action NANONET – Nanomechanics of intermediate filament networks. We gratefully acknowledge Audrey Salles and the Imagopole of Institut Pasteur (Paris), and the financial support of the Institut Pasteur (Paris), the France–BioImaging infrastructure network supported by the French National Research Agency (ANR-10–INSB–04, Investments for the future), and the Région Ile-de-France (program DIM-Malinf) for the use of the Elyra microscope and for funding the development of a dual objective STORM system (‘NANOINF’ project, to CZ). This work was supported by the French National Research Agency: ANR-16-CE13-0019 (to CL), ANR-18-CE13-0026 (to GG), ANR-16-CE92-0034 (to GG), the Ligue contre le cancer EL2017.LNCC (to SEM), the Centre National de la Recherche Scientifique and Institut Pasteur.

## Author contributions

C.L. conceived and coordinated the project. C.L, G.G., S.E.M., C.Z, F.N.V and M.L. designed the research. F.N.V. performed and analysed the experiments combining cell stretching and SRM. M.L. performed and analyzed the experiments with the dual objective microscopy setup. J.Y.T wrote the software for the automatic analysis of pairwise distances along filaments. G.P-A performed EM imaging. C.L. performed and analyzed 2D-dSTORM imaging and cloned the vimentin constructs. DQT purified, labeled vimentin and prepared purified vimentin filaments. C.L and G.G. wrote the manuscript with input from all authors.

## Competing interests

The authors declare no competing interests.

## Data and materials availability

All data needed to evaluate the conclusions in the paper are present in the paper and/or the Supplementary Materials.

## Notes

### Competing Interest Statement

The authors have declared no competing interest.

